# Genome-scale CRISPRi profiling reveals metabolic vulnerabilities of uropathogenic *Escherichia coli* in human urine

**DOI:** 10.1101/2025.10.31.685768

**Authors:** Monica Ortelli, Ruta Prakapaité, Antonia Müller, Solange Miele, Cinzia Fino, Steffi Klimke, Sarah Tschudin-Sutter, Urs Jenal, Carlos Flores, David Bikard, Christoph Dehio

## Abstract

Urinary tract infections (UTIs) are among the most common infectious diseases, causing over 400 million cases and 260,000 deaths annually. Women are disproportionately affected, with ∼50% experiencing at least one UTI during their lifetime and 20–30% suffering from recurrent infections. Uropathogenic *Escherichia coli* (UPEC), which accounts for ∼75% of cases, employs diverse virulence factors to persist and evade host immunity. Rising antibiotic resistance, driven by widespread antimicrobial misuse, is eroding treatment efficacy and highlights the urgent need for alternative therapeutic strategies. To uncover novel vulnerabilities under physiologically relevant conditions, we constructed a genome-wide CRISPR interference (CRISPRi) library in the UPEC reference strain *E. coli* CFT073 and systematically profiled gene fitness in rich media versus human urine. The screen revealed multiple pathways that are conditionally essential for UPEC growth in urine, including iron uptake, envelope maintenance, and the biosynthesis of arginine, methionine, and branched-chain amino acids. Notably, we identified acetolactate synthase (ALS) II as the sole active isoform supporting branched-chain amino acid synthesis in urine. Functional validation further demonstrated its druggability: introducing a re-sensitizing mutation overcame the protein’s intrinsic resistance to the ALS-targeting herbicide sulfometuron methyl, restoring sensitivity. These findings establish ALS II as a promising therapeutic target against UPEC.

## Introduction

Urinary tract infections (UTIs) are among the most common infectious diseases worldwide, with an estimated 405 million cases and 267,000 associated deaths annually^1^. Their incidence varies by age and sex, disproportionately affecting children, the elderly, and young women^2–4^. Epidemiological studies indicate that ⁓50% of adult women experience at least one UTI during their lifetime, and 20-30% suffer from recurrent episodes^4,5^. In healthcare settings, UTIs account for 20-50% of all healthcare-associated infections^6–8^, underscoring their clinical and socioeconomic burden.

UTIs are caused by a range of pathogens, including Gram-negative and Gram-positive bacteria as well as certain fungi, with uropathogenic *Escherichia coli* (UPEC) being the dominant etiological agent, responsible for ⁓75% of cases^9,10^. UPEC harbors a broad arsenal of virulence factors that promote colonization, invasion, persistence, and immune evasion^11^. Compounding this, UPEC strains frequently exhibit high levels of antibiotic resistance, a problem exacerbated by widespread misuse and overuse of antimicrobials^2,3,12,13^. Since antibiotics remain the first-line treatment for UTIs, the escalating resistance crisis represents a major challenge for effective managment^13^, highlighting the urgent need for alternative therapeutic strategies. A deeper understanding of UPEC physiology in the stringent urinary tract microenvironment, together with systematic identification of conditionally essential genes, is critical to uncover novel bacterial vulnerabilities that can be exploited for intervention.

To address this, we constructed a pooled CRISPR interference (CRISPRi) library targeting the entire genome of the UPEC reference strain *E. coli* CFT073^14^. This platform combines a catalytically inactive Cas9 (dCas9) with a genome-wide single-guide RNA (sgRNA) library to enable sequence-specific transcriptional silencing. By monitoring sgRNA abundance before and after growth in nutrient-rich laboratory media and in human urine, we defined gene fitness profiles as a proxy for essentiality^15–17^. Our results aligned well with previous genome-wide analyses of UPEC in urine, reinforcing established metabolic dependencies while revealing additional genetic determinants required for growth under these conditions. In particular, we observed a strong dependence on branched-chain amino acid (BCAA) biosynthesis, and we also identified acetolactate synthase II (ALS II) as the sole ALS isoform active in urine and a possible drug target of this central metabolic pathway.

## Results

### EcoCFT073 CRISPRi library generation and validation

Genome-wide CRISPRi screens have emerged as a powerful approach to assess gene essentiality and functional requirements in bacteria across diverse conditions^18,19^. However, this popular approach has not yet been established for UPEC at whole genome scale. To address this gap, we constructed a pooled CRISPRi library targeting the genome of the UPEC reference strains *E. coli* CFT073 (EcoCFT073 library). Our design built on the previously published *E. coli* core genome library (EcoCG library)^20^, which covers nearly all genes present in >90% of *E. coli* genomes available in Genbank as of 2018. To this core set, we added sgRNAs targeting nearly all additional genes present in *E. coli* CFT073, generating a complementary “CFT073_Add-on” library.

Mapping EcoCG onto the CFT073 genome showed that 10,530 of 11,629 sgRNAs uniquely target a single CFT073 gene (RefSeq entry: NZ_CP051263), covering 64.5% of the genome. By supplementing with CFT073-specific guides, we increased coverage to 97% while maintaining a compact library of 15,173 sgRNAs in total. The final EcoCFT073 library targets 98.1% of protein-coding sequences (CDS), 91.3% of rRNA genes, 42% of tRNA genes and 83% of annotated ncRNA. Each gene is represented by 3-4 sgRNAs to ensure robust measurements of fitness effects (**Supplementary Table 1)**.

To ensure compatibility with EcoCG, all sgRNAs of CFT073_Add-on were cloned into pFR56, a dCas9-sgRNA expression vector optimized to avoid most *E.coli* restriction modification systems and to ensure non-toxic expression of dCas9^20^. The pooled EcoCFT073 library was generated in biological triplicates by adapting the protocol from Rousset et al.^20^ (see *Methods*). This procedure ensured optimal representation, with >99% of designed sgRNA recovered in each replicate and an average per-guide coverage of about ⁓1000-fold (**Extended Data Figure 1a**).

We next benchmarked the library’s accuracy in predicting gene essentiality. Fitness contribution of each gene was quantified by measuring the fold change (FC) in sgRNA abundance during growth in Lysogeny Broth (LB). Consistent with previous studies^20^, strong fitness defects (here any gene with a median log_2_FC ≤-2) were used as proxy for gene essentiality, while strongly increased fitness (median log_2_FC ≥ +2) were considered costly. Notably, 77.8% of the genes absent from the Keio knockout collection, and therefore considered essential^21^ exhibited a strong fitness defect in our screen, confirming the high reliability of the EcoCFT073 library for identifying essential genes (**Extended Data Figure 1b** and **Supplementary Results**).

### Genome-wide CRISPRi reveals diverse conditionally essential genes in rich media and urine

Bacteria have distinct genetic requirements for growth and survival across environments. Defining these condition-specific adaptations is essential for identifying vulnerabilities that may serve as selective antimicrobial targets. Using the EcoCFT073 CRISPRi library, we profiled gene essentiality in two laboratory rich media (LB and MHB) and in pooled human urine collected from healthy donors, monitoring bacterial fitness across ⁓20 bacterial generations (**Figure 1a**).

**Figure 1:**
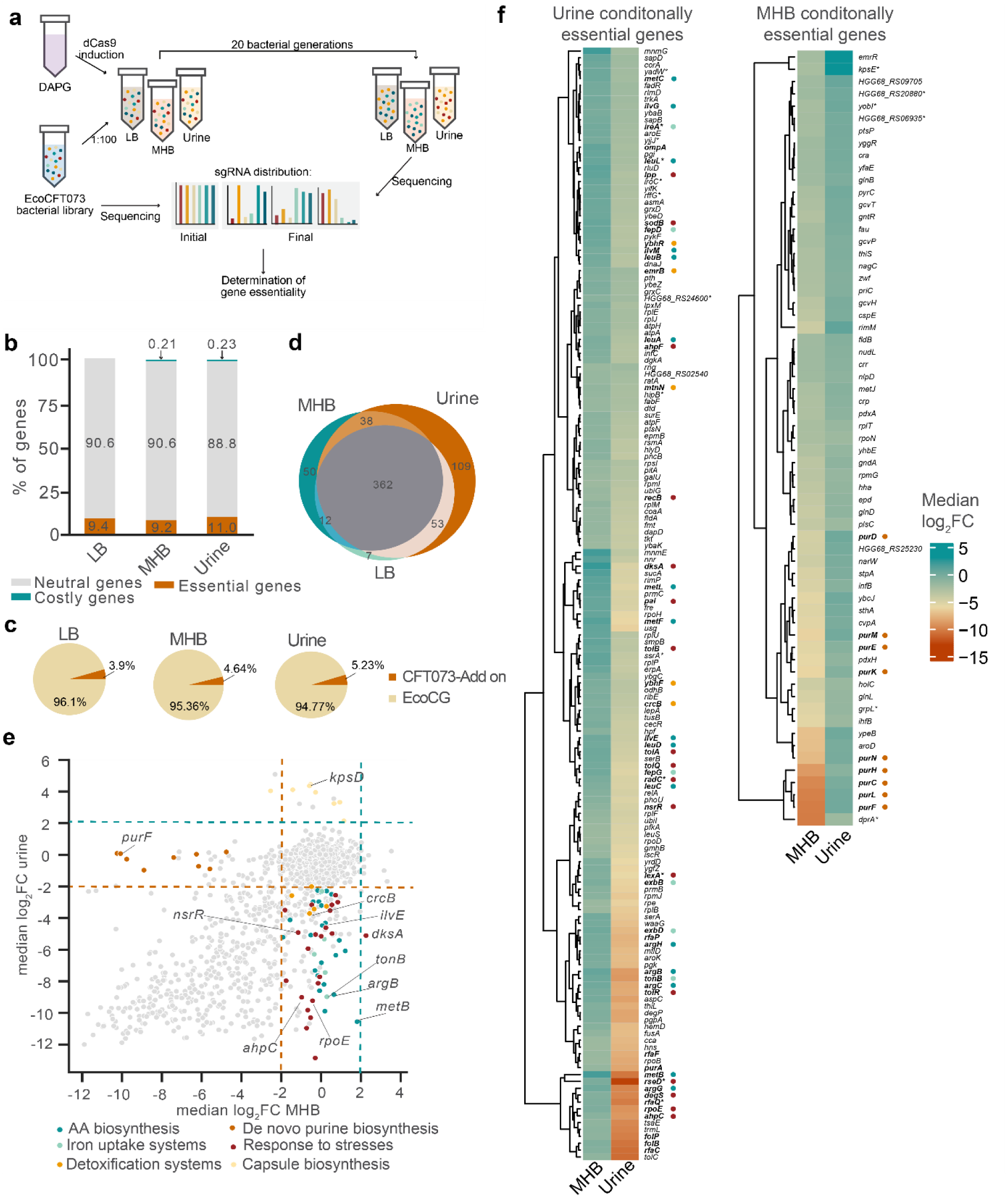
Genome-wide CRISPRi screening in EcoCFT073 for conditionally essential genes in rich media and human urine. **a**, Schematic of the pooled CRISPRi screening workflow. **b**, Distribution of genes classified as neutral, essential, or costly for growth in LB, MHB, or human urine. **c**, **Relative contributions** of the EcoCG and CFT073_Add-on libraries to the conditionally essential gene sets. **d**, Venn diagram showing the overlap between core essential genes (grey) and conditionally essential genes identified in LB, MHB or urine. **e**, Pair-wise comparison of gene essentiality between MHB and urine. Scatter plot depicts median log₂ fold-change (log₂FC) values; conditionally essential genes are color-coded by functional category. A representative gene for each functional category discussed in the main text is labelled in black. **e**, Heat map of median log₂FC values for urine versus MHB across conditionally essential genes. A colored dot next to each gene discussed in the main text indicates its functional category. Genes targeted by the CFT073_Add-on library are marked with an asterisk.

By calculating the median of the log_2_ fold-change (log_2_FC) of each targeted gene, we observed that the number of essential genes varied with growth medium (**Figure 1b**). Roughly 9% of genes were required to grow in rich media, compared to 11% in urine. Genes targeted by the CFT073_add-on library comprise 5.23% of the conditionally essential gene set in urine, compared to 3.90% in LB and 4.64% in MHB, indicating a substantial contribution of the CFT073 accessory genome to the essentiality landscape under different conditions (**Figure 1c**).

Similarly, the fraction of costly genes was medium dependent: 0.21% in MHB, 0.23% in human urine pool and none in LB. The vast majority of genes, 90.6% in LB, 90.5% in MHB and 88.7% in urine, showed no substantial impact on growth (-2 ≤ log_2_FC ≤ 2; **Figure 1b**).

Of the 631 genes required in at least one condition (pan-essential gene set), 57% (362 genes) were essential across all three conditions (core-essential gene set), encoding for core cellular functions such as DNA replication and transcription (**Sup. Table 2**). Considering the entire pan-essential gene set, LB and MHB exhibited high similarity, sharing 86% and 80% of their essential genes, respectively. In contrast, genes required for growth in urine showed greater divergence, with 17% of the pan-essential genes uniquely required in this condition (**Figure 1d, Supplementary table 3**. Among these, 10% were accessory genes targeted by the CFT073_add-on library. These findings highlight that essentiality is strongly environment dependent, and that MHB, commonly used as reference medium in antibiotic research and development, may not capture requirements relevant for growth in urine. We therefore compared gene essentiality between MHB and urine directly (**Figure 1e**).

The urine conditionally essential genes reflected the unique requirements for *E. coli* CFT073 growth in this nutrient-limited environment. These findings are broadly consistent with previous genome-wide studies of UPEC using transposon-insertion sequencing^22–25^, confirming core requirements for growth in urine. However, approximately two-thirds of the genes identified here represent previously unrecognized contributors to fitness, expanding our understanding of the genetic basis of UPEC adaptation to urine (**Supplementary Table 4** and **Supplementary Results**).

To better characterize these urine conditionally essential genes, we grouped them into broad functional categories based on Gene Ontology (GO) biological process annotations (**Supplementary Table 4**). Following this classification, the largest category comprised amino acid biosynthesis genes (15%), including pathways for arginine (*argB*, *argC*, *argG*, *argH*), methionine (*metB*, *metC*, *metF*, *metL*), leucine (*leuL*, *leuA*, *leuB*, *leuC, leuD*) and isoleucine (*ilvE*, *ilvG*, *ilvM*) synthesis, consistent with the scarcity of these amino acids in urine^26^. Another major group (11%) included folate biosynthesis (*ygiG*, *folP*, *metF*), highlighting the reliance on *de novo* folate synthesis^27,28^. Iron limitation was also reflected by essentiality of uptake systems (*fepD*, *fepG*, *tonB, exbB, exbD, ireA*). Interestingly, siderophore biosynthesis genes were not required, likely due to redundancy, whereas among siderophore receptors dependent on TonB-ExbBD system^29^, only *ireA* showed a fitness defect when lost, suggesting its favored role in iron acquisition under these conditions. Corroborating this, *ireA* expression has been reported to increase in urine compared to LB, supporting its importance in this environment.^30^

Bacterial fitness in urine also depended on genes involved in stress response pathways. The alternative σ^E^ factor (*rpoE*) and its modulators (*rseD*, *degS*) were essential, together with σ^E^-regulated envelope maintenance genes (*ompA*, *rfaC*, *rfaF*, *rfaP* and *rfaQ*). Additional genes encoding envelope stabilizers, including *lpp* and the Tol-Pal complex (*tolQ*, *tolR*, *tolA*, *tolB*, *pal*), were also required, further highlighting the need for preserving membrane integrity in this medium. Moreover, the essentiality of genes counteracting oxidative (*ahpC*, *ahpF, sodB*), nitrosative (*nsrR*), and DNA-damage stress (*radC, lexA, recB*, *dksA*), as well as detoxification and efflux genes (*crcB*, *ybhF*, *ybhR*, *mtnN*, *emrB*), further emphasizes the hostile urine nature.

In contrast, conditionally essential genes in MHB reflected distinct functional categories. The largest categories included transcriptional regulation (20.6%), genes of unknown function (19%), de novo purine biosynthesis (*purC*, *purD*, *purE*, *purF*, *purH*, *purK*, *purl*, *purM* and *purN*; 14.2%), amino acid catabolism (4.7%), carbohydrate derivative transport (4.7%) and translation (4.7%) (**Figure 1e, f**). The observation that de novo purine biosynthesis is essential in MHB but dispensable in urine suggests that urine contains purines at sufficient concentrations to support growth. Indeed, in line with previous studies^25^, this interpretation is supported by the essentiality of genes involved in the salvage pathway (*purA*, *purB*, *guaA*, *guaB*, *hpt*), which are also required in MHB (**Figure 1e, f; Supplementary Table 5**).

The contrasting urine and MHB profiles illustrate how *E. coli* CFT073 tailors its genetic program to environmental pressures: urine drives nutrient acquisition and stress resistance, while MHB favors transcriptional control and nucleotide biosynthesis. These findings highlight the limitations of inferring in vivo-relevant physiological states from standard laboratory media, which do not faithfully reproduce the conditions encountered in the urinary tract.

Interestingly, we also identified conditionally costly genes, eight in MHB and twelve in urine. The latter includes eight genes involved in capsule biosynthesis (*kdsB, kpsC, kpsD, kpsE, kpsF, kpsM, kpsS, kpsT*), indicating that capsule biosynthesis occurs in urine and imposes a measurable metabolic cost as already pointed out by others^31^ (**Figure 1e**, **Supplementary Table 6**).

### Validation of BCAA biosynthesis as conditionally essential in human urine

To further investigate the biological relevance of genes uniquely required for growth in urine, we performed functional enrichment analysis of urine conditionally essential genes using STRING (**Supplementary Table 7**). Strikingly, three out of the ten most enriched biological processes were associated with branched-chain amino acid (BCAA) biosynthesis, including the general BCAA biosynthetic process (GO:009082), and specifically the leucine (GO:00909) and valine (GO:0009099) biosynthetic pathways (**Extended Data Figure 2a**). These findings led us to assess the CFT073 BCCA synthesis pathway in vitro.

BCAA biosynthesis proceeds via a branched pathway in which production of valine and isoleucine share several enzymatic steps. The first shared step is catalyzed by acetolactate synthase (ALS), which initiates isoleucine biosynthesis and catalyzes the second step in valine biosynthesis^32^ (**Figure 2a**). *E. coli* encodes three ALS isoenzymes, ALS I (catalytic: *ilvB,* regulatory: *ilvN*), ALS II (catalytic: *ilvG*, regulatory: *ilvM*) and ALS III (catalytic: *ilvI*, regulatory: *ilvH*). While these three enzymes catalyze the same reaction, they differ in feedback regulation: ALS I and ALS III are inhibited by valine, whereas ALS II is resistant to valine inhibition^32,33^.

**Figure 2:**
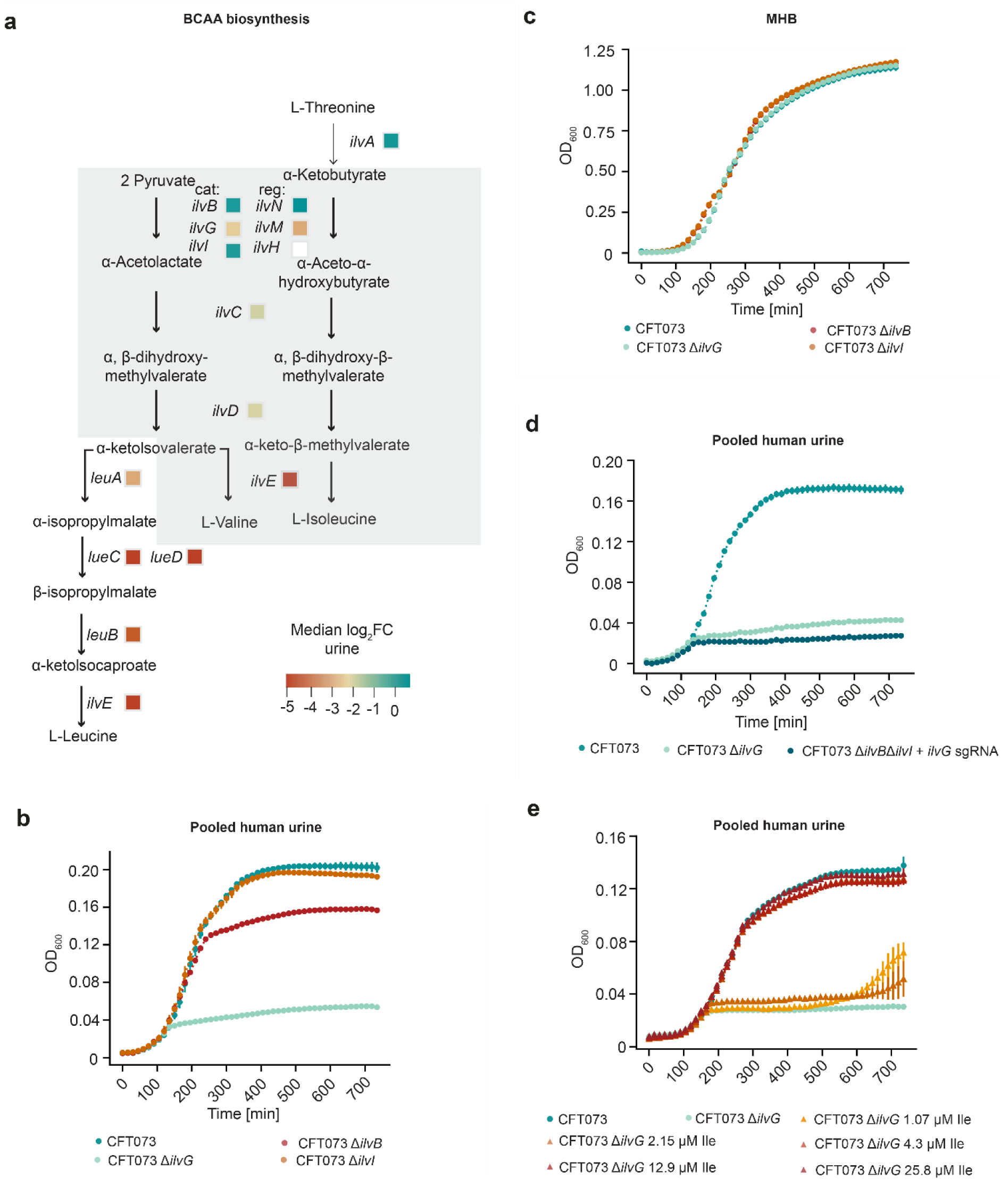
Validation of BCAA biosynthesis as conditionally essential in human urine. a, Schematic representation of the BCAA biosynthetic pathway. Urine median log_2_FC of each gene in this pathway is shown next to the corresponding gene name. The grey box highlights L-valine and L-isoleucine biosynthesis steps which are catalyzed by the same enzymes. Cat: catalytic subunit, reg: regulatory subunit. **b, c,** Growth dynamic comparison of *E. coli* CFT073 wild-type, Δ*ilvB,* Δ*ilvG* and Δ*ilvI* in human urine (c) and MHB (d) over 12 h. **d,** Growth dynamic of *E. coli* CFT073 Δ*ilvB* Δ*ilvI* with CRISPRi-*ilvG* in human urine over 12 h. dCas9 expression was induced supplementing urine with 50 µM DAPG. **e,** *E. coli* CFT073 Δ*ilvG* growth dynamic in dynamic in human urine supplemented with increasing concentration of isoleucine. The growth of CFT073 wild-type and Δ*ilvG* in unsupplemented urine is shown for comparison. All the growth curves represent the mean ± standard deviation of three biological replicates.

CRISPRi profiling indicated that only ALS II was conditionally essential in urine, with *ilvG* repression resulting in a median log_2_FC of-2.25, while targeting ALS I (*ilvB*) or ALS III (*ilvI*) had no significant effect on fitness (median log_2_FC 0.09 and 0.04, respectively) (**Figure 2a** and **Supplementary Table 3**). To validate these results, we constructed CFT073 knockout strains lacking the catalytic subunit of each ALS isoform.

In pooled human urine, the Δ*ilvG* mutant exhibited normal early-phase growth but rapidly entered growth arrest (**Figure 2b**). By contrast, Δ*ilvG* grew comparably to the wild type in MHB, confirming that *ilvG* is specifically required in urine (**Figure 2c**). Growth in urine was unaffected by deletion of *ilvI* and only slightly reduced by deletion of *ilvB* indicating that ALS I and ALS III are dispensable under this condition (**Figure 2b**).

To account for variability in urine composition between individuals^26,34^, we further tested Δ*ilvG* growth in 10 individual urines from healthy volunteers. In all cases, Δ*ilvG* displayed similar growth defect, indicating that *ilvG* conditional essentiality is robust and donor-independent (**Extended Data Figure 2b**). We next investigated why only ALS II is functionally essential in urine. Metabolomic analyses have shown that urine contains valine at higher concentrations than isoleucine (3.4 μM versus 1.3 μM)^26^. Because ALS I and ALS III are feedback inhibited by valine^32,33^, their activity would be suppressed under these conditions, leaving ALS II as the sole functional isoform. Consistent with this model, a mutant lacking ALS I and ALS III catalytic subunits and with an ALS II knock-down (Δ*ilvB* Δ*ilvI* / CRISPRi-*ilvG*) phenocopied Δ*ilvG* growth in urine, confirming that ALS II provides the only active ALS in this environment and its absence triggers valine-induced isoleucine starvation (**Figure 2e**).

We further hypothesized that the Δ*ilvG* growth arrest in urine stems from impaired isoleucine synthesis. Supplementing human urine with increasing concentrations of isoleucine progressively rescued Δ*ilvG* growth, with full restoration first observed at 12.9 μM (**Figure 2e**).

Together, these results validate our CRISPRi screen and demonstrate that ALS II (*ilvG*-encoded) is the sole active ALS isoenzyme in human urine. Its activity is essential to maintain BCAA biosynthesis under these conditions, establishing the pathway as a critical requirement for *E. coli* CFT073 growth in the urinary tract.

### A frameshift mutation in *ilvG* impairs growth of *E. coli* K12 MG1655 in human urine

The widely used laboratory strain *E. coli* K12 MG1655 carries a frame-shift mutation in *ilvG*, rendering ALS II non-functional^35^. Given the essential role of *ilvG* in supporting *E. coli* CFT073 growth in urine, we hypothesized that this defect could account for the poor growth of MG1655 in this environment. Indeed, MG1655 growth in human urine closely mirrored that of *E. coli* CFT073 Δ*ilvG* (**Figure 3a**). Restoring *ilvG* function by reverting the frame-shift to the functional in-frame sequence fully rescued growth, yielding a phenotype comparable to wild-type CFT073 (**Figure 3a**).

**Figure 3:**
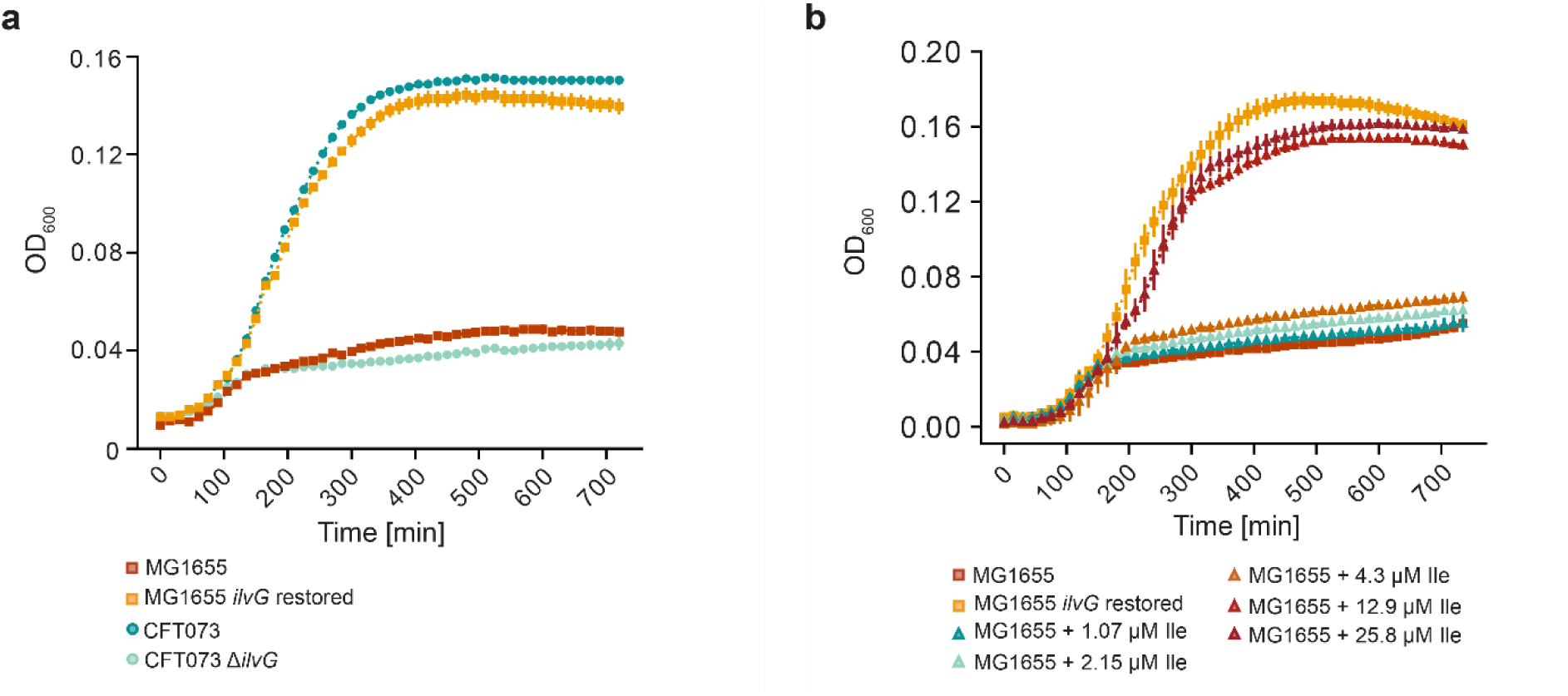
A frameshift mutation in *ilvG* impairs growth of *E. coli* K12 MG1655 in human urine. a, Growth dynamic of *E. coli* CFT073 wild-type, Δ*ilvG*, *E. coli* K12 MG1665 wild-type and a mutated strain with a restored *ilvG* gene in human urine over 12 hours. **b,** Growth curve of *E. coli* K12 MG1665 wild-type in human urine supplemented with increasing concentration of isoleucine. All growth curves represent the mean ± standard deviation of three biological replicates.

As further validation, supplementation of urine with isoleucine progressively rescued MG1655 growth, with full recovery at the lowest effective concentration of 12.9 µM (**Figure 3b**). This effect was identical across pooled and individual donor urine samples (**Extended Data Figure 2c**).

Together, these results demonstrate that the *ilvG* frameshift mutation is the primary factor limiting MG1655 growth in human urine, underscoring the importance of ALS II–mediated BCAA biosynthesis for survival in this niche.

### Role of ALS II during bladder epithelium infection

Having shown genetically that ALS II is required for UPEC growth in human urine (**Figure 2b**), we next investigated whether ALS II contributes to UPEC fitness in host-associated niches beyond urine, where host-derived BCAA might be accessible. To test this, we performed infections in a Transwell-based human urothelial microtissue model^36,37^, quantifying both planktonic and tissue-associated bacteria at early (3 hpi), mid (6 hpi) and late (12 hpi) stages of infection (**Figure 4a**).

**Figure 4:**
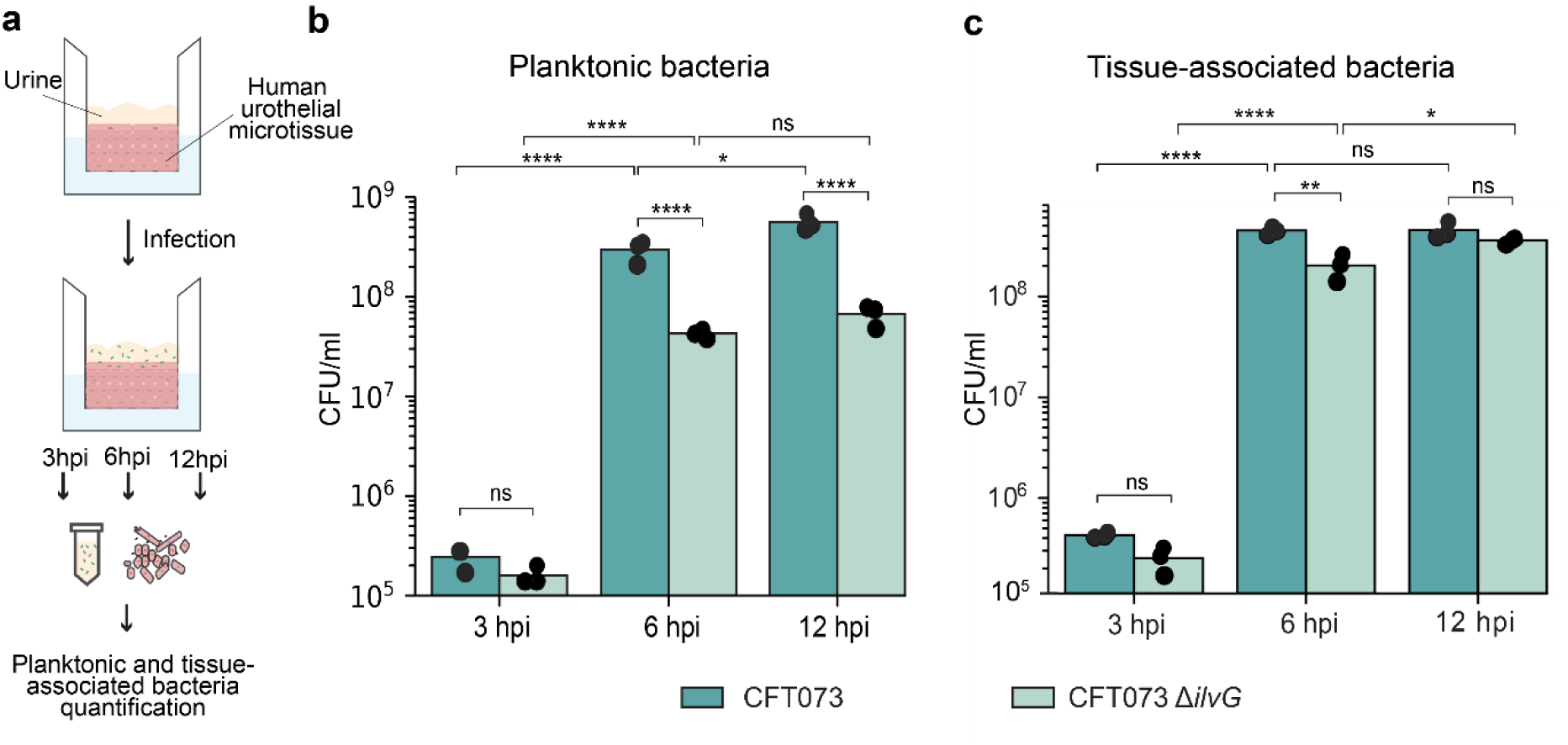
Role of ALS II during bladder epithelium infection. a, Schematic of the experimental setup. **b,** Quantification of planktonic bacteria (CFU/ml) at 3 h, 6 h and 12 h post-infection. **c,** Quantification of tissue-associated bacteria (CFU/ml) at 3 h, 6 h and 12 h post-infection. Each dot represents a biological replicate. Statistical significance was assessed by two-way ANOVA with Tukey’s multiple comparison as post hoc test. Significance levels: ns (p > 0.05), * (p < 0.05), ** (p < 0.01), *** (p < 0.001), **** (p < 0.0001).

For planktonic bacteria, both wild-type and the Δ*ilvG* mutant increased significantly between 3 and 6 hpi (adjusted p < 0.0001). However, while the wild-type strain continued to grow between 6 and 12 hpi, showing a further significant 2.8-fold increase (adjusted p value 0.04), the Δ*ilvG* strain did no significantly grow during this interval. Despite its initial growth, Δ*ilvG* consistently showed ⁓1-log lower CFU/ml than wild-type at both 6 and 12 hpi (adjusted p < 0.0001, **Figure 4b**).

For tissue-associated bacteria, the population of both isogenic strains increased significantly from 3 to 6 hpi (adjusted p < 0.0001). Unlike the planktonic fraction, wild-type levels did not change further between 6 and 12 hpi, suggesting a saturation point of tissue colonization. The Δ*ilvG* mutant, however, showed a slight but significant increase over this interval (adjusted p = 0.03). At 6 hpi, Δ*ilvG* CFU/ml was ∼3.5-fold lower than wild-type, but by 12 hpi this difference was no longer significant, indicating a delayed rather than arrested growth phenotype (**Figure 4c**).

These results suggest that once isoleucine is depleted in the urine compartment, planktonic UPEC requires ALS II for sustained proliferation, consistent with our axenic urine assays. In contrast, upon colonization of the urothelial tissue, the Δ*ilvG* mutant continues to grow slowly, possibly by accessing host-derived isoleucine that partially compensate for the loss of ALS II activity. This host-proximal growth likely explains why tissue-associated bacterial loads exceed those of the planktonic fraction despite the mutant’s reduced growth rate compared to wild-type.

### Chemical targeting of ALS II in UPEC

Because ALS II is important for UPEC growth in urine, as demonstrated both in axenic conditions and during infection (**Figures 2b and 4b)**, we next tested whether this enzyme can be targeted in this environment by small molecular inhibitors. Because the BCAA biosynthesis pathway is absent in animals but conserved in bacteria, fungi, and plants, it has long been considered an attractive selective target^33,38–41^. For instance, in agriculture, ALS enzymes are the molecular targets of several widely used antimetabolite herbicides^42,43^. Among them, the sulfonylurea sulfometuron methyl (Sm) inhibits ALS II and ALS III, but not ALS I^44^. Since ALS II is the sole active isoform in urine (**Figure 2d**), we used Sm as a probe compound to test whether its inhibition would suppress *E. coli* CFT073 growth in human urine.

In wild-type CFT073, Sm reduced growth in a dose-dependent manner but did not recapitulate the growth arrest of the Δ*ilvG* mutant (**Figure 5a**). Minimum inhibitory concentration (MIC) testing in urine confirmed that growth occurred at all concentrations (**Supplementary Table 8**), indicating intrinsic resistance (MIC > 264 μg/ml). This is probably due to the proline-to-serine substitution at position 100 (P100S) of CFT073 ALSII compared to the *S. cerevisiae* homologue, a mutation previously reported to confer class-wide resistance to sulfonylureas by disrupting binding to the S-ring^45,46^ (**Figure 4b, c**). Mutating serine 100 to proline (S100P) resulted in susceptibility, with a MIC as low as 8.2 μg/ml in urine (**Figure 5d**). Importantly, the sensitized strain grew normally in MHB at the same Sm concentration, reinforcing that ALS II is conditionally essential in urine but dispensable in nutrient-rich medium (**Extended Data Figure 3a**).

**Figure 5:**
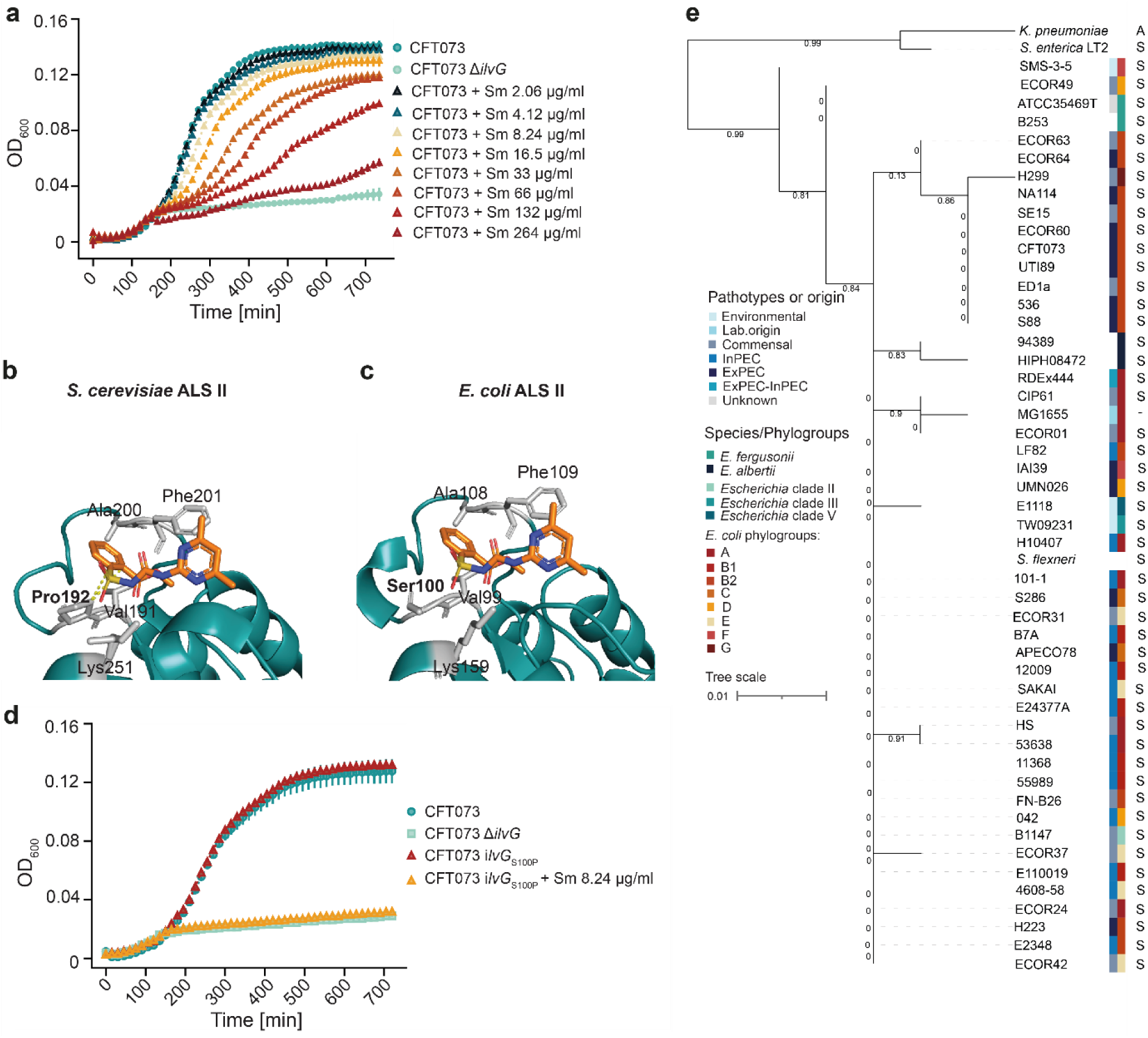
Chemical targeting of ALS II in UPEC. a, Growth curve of *E. coli* CFT073 in human urine supplemented with increasing concentrations of sulfonylurea sulfometuron methyl (Sm). **b,** Structure of *S. cerevisiae* ALS II catalytic subunit bound to Sm (PDB 1T9C^46^). Dashed yellow lines indicate the interaction between P192 and Sm. **c,** Alphafold model of *E. coli* ALS II catalytic subunit^47,48^ with Sm modeled in the ligand-binding pocket, illustrating the missing interaction between Sm S-ring and Ser100. In both structures residues interacting with Sm are shown in grey. **d**, Growth dynamics of *E. coli* CFT073 wild-type and *ilvG_S100P_* mutant in human urine supplemented with 8.2 μg/ml Sm over 12 h. **e**, Phylogenetic analysis of ALS II catalytic subunits of the *Escherichia* genus and closely related *Enterobacteriaceae*. The strains have been selected to represent the phylogenetic diversity of the genus^49^. The tree was reconstructed from multiple protein alignment of ALS II catalytic subunit using PhyML and rooted on *Klebsiella pneumoniae* as an outgroup.

To verify target specificity in the sensitized strain, we supplemented isoleucine in the presence of inhibitory concentrations of Sm. Growth was rescued in a dose-dependent manner (**Extended Data Fig. 3b**), confirming that Sm inhibition is mediated via ALS II rather than by nonspecific toxicity.

Finally, analysis of *Escherichia* genomes from different origins and pathotypes revealed that S100 in ALS II is present in all strains, as well as closely related *Enterobacteriaceae* such as *Shigella flexneri* and *Salmonella enterica* (**Figure 5e and Extended Data Figure 3c**). This indicates that sulfonylurea resistance is a conserved feature within the genus *Escherichia*. Although current herbicide scaffolds are therefore unlikely to be directly transferable, our results identify ALS II representing a vulnerable metabolic node in urine and, in principle, a druggable target.

## Discussion

Understanding the genetic basis of UPEC pathogenesis is critical for developing new therapeutic strategies, particularly as antibiotic resistance continues to rise. To define UPEC requirements during early infection, we constructed the first genome-wide CRISPRi library for the reference strain *E. coli* CFT073 and applied it to high-coverage screens in both human urine and standard laboratory medium (MHB).

The EcoCFT073 library achieves broad genome coverage with compact size and reliably identifies essential genes, as shown by strong concordance with the Keio collection. Unlike transposon mutagenesis, CRISPRi enables inducible and tunable knockdowns, allowing a more precise assessment of gene essentiality^17,50^. This higher resolution revealed both core and pan-essential genes as well as a greater number of urines conditionally essential genes than previous studies. The library’s compact design further facilitates application in settings where transposon libraries face bottlenecks^50–52^, including organoids, 3D tissue, or murine models, while its combinatorial design strategy offers a cost-effective framework readily adaptable to additional UPEC strains.

Our genome-wide screens revealed that essentiality in *E. coli* CFT073 is highly dependent on the growth environment. A conserved set of core functions was required across all conditions, but numerous medium-specific dependencies emerged. In urine, conditionally essential genes reflected two major pressures: nutrient limitation (e.g., amino acid biosynthesis) and host-imposed stresses (e.g., envelope stress responses). These findings were consistent with prior genome-scale analysis of UPEC fitness in urine, validating our approach^22–25^, while also uncovering new contributors such as for example TonB-ExbBD-mediated iron uptake via IreA, and genes involved in oxidative stress response and detoxification. By contrast, MHB yielded a distinct profile, including reliance on both de novo purine biosynthesis and salvage pathways, whereas only salvage was essential in urine.

Notably, de novo purine biosynthesis has been proposed as a drug target in other bacterial pathogens^53–55^, but its apparent essentiality in UPEC grown in standard laboratory media does not extend to host-like environments. This discrepancy underscores how reliance on standard laboratory media can misrepresent bacterial vulnerabilities: pathways dispensable in vivo may appear essential in vitro, while true host-specific requirements may be overlooked. Such biases are especially problematic for transposon-based approaches, where mutants essential in laboratory media are systematically lost during library construction. By contrast, CRISPRi enables systematic, conditional knockdowns without this bottleneck, providing a more faithful picture of context-specific gene requirements. Together, these findings highlight both the advantages of genome-scale CRISPRi screening and the necessity of testing candidate targets under physiologically relevant conditions to ensure that antimicrobial discovery focuses on pathways genuinely required during infection.

As a follow-up of our screening, we examined the branched-chain amino acid (BCAA) biosynthesis. We explored further the long-recognized UPEC dependence on BCAA biosynthesis in urine^25,56–58^, focusing on an early step, catalyzed by acetolactate synthase (ALS), an enzyme that was newly identified as conditionally essential in urine by our screening. This analysis provides mechanistic insight into UPEC dependency: ALS II emerged as the sole active isoenzyme under urinary microenvironment, as high valine concentrations suppress ALS I and III. Consequently, Δ*ilvG* mutants were able to initiate growth but were arrested once the limited isoleucine pool was depleted, a defect that was rescued by isoleucine supplementation. These findings illustrate how conditional essentiality arises directly from the host metabolic environment.

We further showed that the poor growth of the laboratory strain *E. coli* K12 MG1655 in urine is attributable to a frameshift in *ilvG*. Restoring ALS II activity rescued growth, yielding a non-virulent yet genetically tractable model strain for dissecting UPEC physiology under host-like conditions.

Using a bladder epithelial infection model, we uncovered niche-dependent essentiality of ALS II. The enzyme was required for sustained growth of planktonic bacteria but dispensable once bacteria colonized the tissue. Close contact with host epithelium may allow adherent bacteria to access host-derived isoleucine, potentially through localized tissue damage, while invasive bacteria may obtain isoleucine directly from intracellular pools, compensating for the loss of ALS II. In addition, the formation of biofilm-like aggregates on the tissue surface at later stages of infection^37^ could shield tissue-associated bacteria from urine, alleviating valine-induced isoleucine starvation. This dynamic essentiality highlights ALS II as a conditional vulnerability that could be exploited therapeutically only when bacteria remain planktonic, highlighting the potential for synergies with compounds interfering with tissue adhesion^59,60^.

To assess therapeutic potential interfering with ALS, we tested sulfometuron methyl (Sm), a sulfonylurea herbicide known to inhibit ALS II/III^44^. Sm reduced CFT073 growth in a dose-dependent manner, with resistance mediated by a conserved P100S substitution in ALS II. Reversion to proline restored sensitivity, confirming ALS II as the target and showing that inhibition was revealed only under urine conditions, where ALS II is essential. Although sulfonylureas are not clinically viable, they highlight ALS II as a vulnerable metabolic node in UPEC and provide a rational for developing UPEC-specific inhibitors. By analogy to established antimetabolites, such as sulfamethoxazole and trimethoprim, dual inhibition of ALS II and a second BCAA biosynthesis enzyme may yield synergistic activity and limit resistance development. When combined with anti-adhesin strategies that maintain bacteria in the planktonic state^60^, ALS II targeting could maximize efficacy while reducing escape through tissue colonization.

In summary, our study demonstrates that UPEC essentiality is highly context-dependent, with physiologically relevant environments such as urine revealing vulnerabilities that are obscured in standard media. By integrating genome-wide CRISPRi screening with targeted validation, we provide mechanistic insights into conditional essentiality and establish a broadly adaptable platform for antimicrobial target discovery. More generally, our findings illustrate how context-specific genetic requirements can inform rational antimicrobial discovery^61^ and deepen our understanding of UPEC adaptation in the urinary tract.

## Methods

### Urine sample collection and handling

Human urine samples were donated anonymously and separated by gender by healthy volunteers. Donated urine samples were tested using Combur dipstick test (Roche). Samples with anomalous values or a volume less than 50 ml were discarded. Individual urine samples were then either sterile filtered and stocked at-20°C or pooled maintaining an equal number of samples donated from men and women, sterile filtered and stored at-20°C. When needed, a urine sample was thawed at 4°C overnight and then incubated in a water bath at 37°C to dissolve any precipitate.

The anonymous collection of urine samples did not require ethical approval, as confirmed by the responsible ethics committee of north-west and central Switzerland (EKNZ, submission Req-2020-01209).

### Bacterial culture

Unless otherwise stated, bacteria were grown in the growth medium of choice (either LB, MHB or human urine) at 37°C while shaking. LB + 1.5% agar was used as solid medium. If needed, growth media were supplemented with antibiotics and other supplements at the following concentrations: chloramphenicol (Cm) 20 µg/ml, ampicillin (Ap) 30 µg/ml and diamino pimelic acid (DAP). 2,4-diacetylphloroglucinol (DAPG) at a concentration of 50 µg/ml was used to induce dCas9 expression. Induction of lambda red system for gene deletion with 100 µM Isopropyl β-D-1-thiogalactopyranoside (IPTG).

### EcoCFT073 Library design

EcoCFT073 sgRNA library was designed to provide genome-wide coverage of *E. coli* CFT073 (RefSeq entry: NZ_CP051263). The library was generated by combining EcoCG library^20^ which targets *E. coli spp*. core genome, with an additional set sgRNAs specifically designed to target all CFT073 genes not covered by EcoCG (CFT073-specific sgRNAs).

First, we mapped EcoCG library on *E. coli* CFT073 genome to identify CFT073 genes not targeted by EcoCG library and EcoCG sgRNAs uniquely targeted a single CFT073 gene. Then, CFT073-specific sgRNAs were designed as follow: First, we identified all possible sgRNAs located within *E. coli* CFT073 coding regions containing the protospacer adjacent motif (PAM) NGG using generate_library.py^62^. These sgRNAs were then ranked according to the following priority criteria defined by Calvo-Villamañan et al.^62^:

(i) sgRNAs with a unique *E. coli* CFT073 target
(ii) Minimal number of off-targets with perfect 12-nt seed matches (noff_12)
(iii) Minimal number of off-targets on the non-template strand with 11-nt seed matches (noff_11_gene),
(iv) Minimal number of off-targets in the promoter regions with 9-nt seed matches (noff_9_prpmoter)
(v) Absence of 5 nt seed sequences known to silence the promoter of essential genes^63^ (badseed sequences, GGTTA, ACCCA, AGGGG, GATAT)
(vi) Preferential targeting of the first half of the gene
(vii) High activity score predicted by the linear model of Calvo-Villamañan et al.^62^

Based on this ranking, the 4 best sgRNAs targeting each uncovered CFT073 gene were selected, resulting in 4877 CFT073-specific sgRNAs. For cloning into pFR56 plasmid via Golden Gate assembly, each sgRNA was flanked by a specific prefix (AGCTGTCACAGGTCTCATAGT) and suffix containing BsaI (GTTTTGAGACCGCGACTACGTCTAC) recognition sites.

For downstream analysis, approximately 500 EcoCG sgRNAs that do not target any CFT073 gene were used as control guides. Conversely, ∼100 EcoCG sgRNAs with multiple CFT073 targets were excluded due to potential off-targets effects. A complete list of sgRNAs included in the library and considered for analysis is provided in **Supplementary Table 1**.

### CFT073-specific plasmid library generation

CFT073-specific sgRNAs were synthesized as single-stranded DNA oligonucleotides using on-chip oligo synthesis (Twist Bioscience). To ensure high-fidelity and minimize amplification bias, the pooled oligos were amplified using KAPA HiFi DNA polymerase with primer pair Sol71-Sol72 (Thermal cycling conditions: Initial denaturation 3 min at 95°C, 14 cycles of denaturation 20 sec at 98°C, annealing 15 sec at 57°C and extension 15 sec at 72°C, followed by a final extension step at 72°C for 1 min). The amplified oligos were purified according to Nucleo Spin PCR clean up (Macherey-Nagel). The Purified oligos were assembled into the plasmid vector pFR56-ccdB via Golden Gate assembly. The restriction enzyme BsaI v2 (New England Biolabs) was used to digest the plasmid vector. Specific sgRNA targeting *rpsL* and *ilvG* were cloned on pFR56 using the same strategy.

### EcoCFT073 bacterial library construction

To minimize sgRNA loss caused by transient dCas9 expression during cloning, the CFT073-specific plasmid library was electroporated into *E. coli* MG1655::acrII4a (CH-E02), a strain that constitutively expresses the anti-Cas9 protein AcrII4a. To achieve optimal sgRNA-fold coverage, we performed and pooled together 6 electroporations, each with 5 µl of plasmid library and 40 µl of electrocompetent CH-E02 cells. Following 1 h incubation at 37°C, the transformed cells were plated on 12 square LB agar plates supplemented with Cm (500 µl transformant suspension per plate) and incubated at room temperature overnight. The following day, colonies were recovered by washing each plate twice with 1 ml LB with. All the washes were pooled to generate a comprehensive plasmid population. To construct EcoCFT073 plasmid library, plasmids were extracted from the pooled CH-E02 culture by Miniprep and mixed with EcoCG plasmid library balancing both the size and the sgRNA-fold coverage of each library.

To maximize the transfer EcoCFT073 plasmid library into *E. coli* CFT073 we first compared transformation and conjugation efficiency into *E. coli* CFT073 using a test plasmid carrying a sgRNA targeting the gene *rpsL*. As conjugation proved more efficient, the EcoCFT073 plasmid library was electroporated into the DAP auxotroph strain *E. coli* MFDpir. Ten electroporations were performed, each with 5 µl of bacterial library and 40 µl of electrocompetent *E.coli* MFDpir cells, and the resulting cultures were pooled. Because at a temperature up to 42°C dCas9-sgRNA complex formation is inhibited^64,65^, we incubated the transformed MFDpir cells at 42°C for 1 hour to prevent sgRNA depletion. Bacterial cells were then plated on LB agar square plates supplemented with Cm and DAP and incubated at 42°C until colonies were formed. Colonies were then recovered from plates and stored as aliquots at-80°C.

For conjugation, equal volumes of recipient (*E. coli* CFT073) and donor (*E. coli* MFDpir carrying the plasmid library) strains were mixed in LB supplemented with DAP and incubated for 2 h at 42°C. After mating, the mixture was then then plated on square LB agar plates supplemented with Cm and incubated at room temperature overnight. The day after, nascent colonies were harvested from plates and stocked at-80°C. Three independent conjugation experiments were performed to generate biological triplicates of EcoCFT073 bacterial library.

### Genome-wide CRISPRi screening design

To initiate CRISPRi screening, 150 μl of each EcoCFT073 bacterial library biological replicate was inoculated into 1350 μl LB supplemented with Cm and incubated overnight at 37°C with shaking. The day after, cultures were washed twice with PBS to remove traces of growth culture medium. Then, 15 μl of washed culture was inoculated into 1485 μl of fresh medium (LB, MHB or human urine pool) supplemented with Cm and DAPG at proper concentration. The remaining overnight cultures were pelleted and plasmids were extracted to determine sgRNA library composition prior the screen (control samples).

CRISPRi screens were carried out for a total of 20 bacterial generations at 37°C with shaking. To determine the number of passages required to reach 20 bacterial generations, we first calculated the doubling time and number of generations per passage (calculated as exponential phase duration divided by doubling time) in each medium using growth curves and semi-logarithmic analysis. Based on these values we carried out four passages every 2 h 20 min in LB, four passages every 2 h 40 min in MHB and five passages every 3 h 40 min in pooled human urine. Plasmids were then extracted to quantify the sgRNA abundance and library composition in the different media.

### Illumina sample preparation and sequencing

To quantify sgRNA abundance and assess library composition, plasmids were extracted from EcoCFT073 library both before and after screening. Samples were prepared for Illumina sequencing as described below. To enable multiplexing and reduce sequencing costs, we used Illumina Unique Dual Indexing strategy, involving two sequential PCR reactions: Illumina adapter PCR and indexing PCR.

Because regions outside the sgRNA target in the amplified libraries exhibit low sequence diversity, which can reduce Illumina sequencing quality, we designed four different primer pairs for the adapter PCR (**Supplementary Table 9**). 100 ng of each sample was amplified using KAPA HiFi DNA polymerase with one of these primer pairs, which differ by a few bases immediately upstream the adapter sequence increasing thereby sequence diversity. This strategy improved overall sequencing yield. The following thermal cycling conditions were used: Initial denaturation 3 min at 95°C, 25 cycles of denaturation 20 sec at 28°C, annealing 15 sec at 60°C and extension 20 sec at 72°C, followed by final extension 5 min at 72°C. PCR products were then purified using AMPure XP beads (Beckman Coulter) and their quality was assessed with Bioanalyzer DNA 100 (expected PCR product size 278-284 bp depending on the primer pair used).

Successively, we performed index PCR using Illumina DNA/RNA UD indexes. To allow multiplexing during sequencing, each sample was amplified using KAPA HiFi DNA polymerase with a specific index combination as recommended in the index pooling guidelines bulletin on the Illumina website. The following thermal cycler conditions were used: Initial denaturation 3 min 95°C, 8 cycles of denaturation 30 sec at 95°C, annealing 30 sec at 55°C and extension 30 sec 72°C, followed by final extension of 5 min at 72°C. Afterwards, PCR products were purified with AMPure XP beads and checked for size, concentration and purity with Bioanalyzer DNA 100(expected PCR product size 351-357 bp depending on the primer pair used while performing Illumina adapter PCR).

Samples were pooled and sequenced with NextSeq 500 sequencer using Illumina NextSeq500/550 high output kit v2.5 (75 cycles) performing 25 cycles of single-end sequencing and 10 cycles to read each index. We obtained an average of ∼217 million read per sample, which correspond to a coverage of ∼ 800x.

### CRISPRi screening data analysis

Following sequencing, indexes sequences were used to perform demultiplexing directly on the Illumina NextSeq500 machine with local run manager software. Since Illumina NextSeq500 produces four different FASTQ files per sample (one per lane), files corresponding to the same condition were merged to generate a single FASTQ file per sample. CRISPRi screens were performed in biological triplicates, and all three replicates were included in downstream analysis.

The quantification of sgRNA abundance was performed using FAST2Q software^50^ with the following parameters:

(i) Feature start position in the reads. This indicates the first nucleotide of the sgRNA within the read, which depends on the Illumina adapter PCR primers used. Specifically, sgRNA start at position25 for primers prMO028-29, 26 for prMO030-31, 27 for prMO032-33 and 28 for prMO034-35.
(ii) Feature length: 20 bp.
(iii) Minimal feature Phred score: 30, corresponding to a base call accuracy of 99.9%.

Low abundance sgRNAs with less than 20 total reads across all replicates were filtered out (about 0.6 % of the library). To determine the enrichment or depletion of the remaining sgRNAs, we then calculated the log2FC between the induced and non-induced conditions using the R package DESeq2^66^. DESeq2 incorporates the three biological replicates to estimate variance across samples, allowing the calculation of adjusted p-values for each sgRNA and providing statistical confidence in the measured effect. Non-targeting control sgRNAs were used for normalization as they should not have any effect on bacterial growth.

We then determined the fitness effect at the gene level by calculating the median log2FC of all sgRNAs targeting the same gene. A median log2FC ≤-2, corresponding to a ≥ 75% reduction in abundance, was chosen as threshold for identifying genes with a strong negative impact on bacterial growth. This relaxed threshold allowed the detection of both genes causing major fitness defects and fully essential genes, which typically reach a median log2FC ≤-4.4, as imposed by sequencing depth limit (log_2_(1/21) ≈-4.4). For simplicity, throughout this study we refer to genes with a median log2FC ≤-2 as essential or conditionally essential, acknowledging that this category includes genes with major fitness defects that are not strictly lethal. Conversely, genes with a median log2FC ≥ + 2, corresponding to a 300% increase in abundance, were classified as costly for bacterial growth.

### Gene deletion and gene insertion with λ-red recombination system

Deletion and insertions mutants were constructed using λ-red recombination system. A double selectable cassette (DScas) encoding chloramphenicol resistance gene (*cat*) and *sacB* for counter-selection was amplified with primers having 50 base overhangs homologous to the flanking regions of the target gene.

To induce λ-red recombinase expression into *E. coli* CFT073, cells were transformed with plasmid pKM208^67^ and grown in LB at 30°C. When cultures reached an *E. coli* CFT073 an OD_600_ of about 0.5 was reached, the expression of λ-red recombinase was induced with 1 mM IPTG for 1 hour. Successively, cells were made electrocompetent and electroporated with purified *cat-sacB* cassette. Recombinants were selected on LB agar plates supplemented with Ap and Cm at proper concentration and *cat-sacB* cassette integration was verified by colony PCR.

For the generation of deletion mutants, a single strand oligonucleotide bridging the upstream and downstream flanking region of the target gene was electroporated into strains carrying the integrated double-selectable cassette and expressing λ-red recombinase. Fog gene insertion, the gene of interest was amplified with primers carrying 50 bases overhang homologous to the insertion site and then electroporated under the same conditions. After electroporation, cultures were incubated for 3 hours to allow recombination and plated on LB agar plate supplemented with Ap and 20% sucrose to select for cassette loss via *sacB*-mediated counter-selection. Finally, plasmid pKM208 was cured and successful gene deletion or insertion was verified by Sanger sequencing.

All the primers and single strand oligos used to generate deletion and insertions mutant are listed in **Supplementary Table 9**.

### Creation of *E. coli* CFT073 *ilvG*_S100P_

The S100P point mutation was introduced into *E. coli* CFT073 *ilvG* gene using overlap extension PCR (OE-PCR). The gene was first amplified in two separate fragments with complementary overlapping sequences at their 3’ and 5’, respectively. The first fragment covered the 5’ portion of the gene up to the mutation site and was amplified with primers pair prMO241-242, with prMO242 carrying a 20 pb overhang encoding the desired point mutation. The second fragment spanned the remaining portion of the gene was amplified using primers pair prMO243-244, with primer prMO244 containing 20 bp overhang complementary to that of primer prMO242. The two fragments were amplified by PCR using KOD DNA polymerase with the following thermal cycler conditions: initial denaturation 2 min at 95°C followed by 30 cycles of denaturation 15 sec at 95°C, annealing 5 sec at 60°C and extension 10 sec at 68°C.

The two PRC products were purified (Gen clean up kit Promega) and mixed at a 1:1 molar ratio. To allow the annealing and extension across the overlapping regions we performed 15 cycles of denaturation 15 sec at 95°C, annealing 5 sec at 60°C and extension 10 sec at 68°C. Subsequently, primer pair prMO241-prMO244 were added to the reaction and an additional 20 cycles were performed to amplify the joined product (denaturation 15 sec at 98°C, annealing 5 sec 60°C and extension 10 sec at 68°C). The final PCR product was purified and Sanger sequenced to verify the presence of the desired mutation.

To insert *ilvG*_S100P_ into *E. coli* CFT073 chromosome we amplified it with primer pair prMO245-prMO246, which carry 50 bp overhang complementary to the upstream and downstream region of the insertion site. We then inserted this gene into *E. coli* CFT073 *ilvG*::DScas performing λ-red recombineering as described in the previous section.

### Growth curves

A single colony of each desired strain was picked from LB agar plate and inoculated into LB overnight at 37°C, with shaking. The next day, overnight cultures were washed twice with PBS and diluted to an OD_600_ of 0.01 into the appropriate growth medium. If necessary, growth media were supplemented with antibiotics or other additives. Growth dynamics were assessed using an automated plate reader (Synergy H4 Hybrid reader, Biotek), which recorded OD_600_ every 15 minutes over a 12 h period while continuously shaking the plate. For each condition, growth curves were measured in three independent biological replicates.

### MIC determination

Sulfometuron-methyl (Sm) susceptibility was essayed performing microdilution MIC testing both in cation adjusted MHB and pooled human urine following the procedure described in Broth microdilution-EUCAST reading guide v 5.0. Bacterial cultures were adjusted to 5 x 10^5^ CFU/ml and incubated at 37°C for 18-20 hours under static conditions. The following Sm concentrations were tested: 2.06, 4.12, 8.25, 16.5, 33, 66, 132, 264, 524, 1056, 2112 µg/ml.

### Modeling of *E. coli* ALS II catalytic subunit interaction with sulfometuron-methyl (Sm)

Because the crystal structure of the *E. coli* ALS II catalytic subunits has not been resolved yet, we used the AlphaFold-predicted model (AF-P0DP90-F1-v4) ^47,48^. The predicted structure was superimposed onto the crystal structure of the yeast ALS II catalytic subunit bound to Sm (PDB 1T9C^46^) using PyMOL (Schrödinger, LLC) to visualize and highlight the putative Sm-binding region in the *E. coli* model.

### Phylogenetic analysis of ALS II catalytic subunit of *Escherichia* genus

IlvG protein sequences were extracted from a set of strains selected to represent the phylogenetic diversity of the Escherichia genus^49^ (**Supplementary Table 10**). Sequences were aligned using R package msa^68^ and the resulting alignment was used to construct a maximum-likelihood phylogenetic tree with the Phylgeny.fr web tool in the “one click” mode^69^. The phylogenetic tree was visualized and annotated using iTOL version 6^70^.

### Generation of human urothelial tissue model

Transwell-based 3D human urothelial models were generated as previously described^36,37^, with a slight modification. Briefly, cells were seeded onto 6.5 mm, 0.4 µm pore polycarbonate filter membranes in plastic inserts standing in 24-well Transwell plates (Corning) in CnT-Prime medium, and after proliferation, they were exposed to barrier medium (CnT-Prime-3D medium, CELLnTEC) from the basal chamber and human pooled urine from the apical chamber.

### Tissue infection and detachment assay

Two days prior to infection, a single colony of *E. coli* CFT073 WT and *E. coli* CFT073 Δ*ilvG* was inoculated into LB and grown statically at 37°C for 48 hours. Cultures were washed twice with pooled human urine and bacterial concentrations were quantified using QUANTOM Tx Microbial Cell Counter (Logos) according to manufacturer’s instruction. For infection, bacteria were inoculated at 1 x 10^5^ cells/ml in the apical chamber in pooled human urine and incubated for 3, 6 and 12 h at 37°C, 5% CO_2_. A 3 h infection correspond to an early infection stage, where *E.coli* CFT073 is mainly scattered on the apical urothelial surface a low density; 6 h to a mid-stage of infection, with bacteria starting to form aggregates on the surface and invading the tissue, while 12 h correspond to a later stage of infection with substantial invasion and possible intracellular bacterial community (IBC) formation^37^.

At each infection time point, apical milieu was collected, the tissues were washed three times with 60 μl 1x PBS and the washed out was pooled with the apical content to retrieve the entire planktonic bacterial fraction. This fraction was then serially diluted and plated on square LB agar square plates for CFU quantification. To assess tissue-associated bacteria, 100 μl of 1% triton X-100 was added to the apical side of each insert (without the apical milieu) and incubated for 20 min at 37°C, 5% CO_2_. Tissues were then lysed by mechanical disruption, scraping the surface of the membrane. The resulting lysates were collected, serially diluted and plated for CFU determination. Each infection condition was performed in biological triplicates.

### Statistical analysis

Statistical analysis was performed using python. CFU counts from bladder epithelium infections were log-transformed prior to analysis, then to assess whether there is significant difference between strains and over time, two-way ANOVA was performed. Tukey’s multiple comparison was used as post-hoc test. Significance levels were reported as follows: ns (p > 0.05), * (p < 0.05), ** (p < 0.01), *** (p < 0.001), **** (p < 0.0001).

## Data availability

CRISPRi raw sequencing data have been deposited in the European Nucleotide Arichive (ENA) under study accession PRJEB101799.

## Code availability

CFT073_Add-on CRISPRi library was generated using “EcoCFT073 add on library design.ipynb”.

## Author contributions

M.O., C.F. (Cinzia Fino), D.B. and C.D. conceived and designed the study. M.O. constructed the CFT073 guide RNA library with support from S.M. and D.B., and performed all the experiments, with input from R.P. and U.J. for bacterial genetics and C.F. (Carlos Flores) for urothelial generation and infection. M.O. analyzed and interpreted the results and drafted the manuscript with input from all co-authors. A.M. provided technical help. S.K. and S.T. provided support for urine collection and ethical consideration. D.B. and C.D. provided intellectual input. All authors read and approved the final manuscript.

## Supporting information

Extended Data Figures and supplementary results

Python script used to generate EcoCFT073 CRISPRi library

## Acknowledgment

We thank all members of the Genomics Facility Basel for their support in sequencing, as well as the members of the Dehio and Bikard teams for valuable technical and scientific discussions. We are also grateful to Dr. Julie Sollier for her insightful input and support. This work was funded by the Swiss National Science Foundation NCCR AntiResist (51NF40_180541) to C.D., the European Research Council (grant agreement no. 101044479) to D.B., and the Agence Nationale de la Recherche (ANR-10-LABX-62-IBEID) to D.B., C.F. acknowledges the support by the International Human Frontier Science Program Organization (HFSPO), under the long-term fellowship LT0017/2023-L.

## References

1. Yang, X. et al. Disease burden and long-term trends of urinary tract infections: A worldwide report. Front. Public Health 10, 888205 (2022).

2. Foxman, B. The epidemiology of urinary tract infection. Nat. Rev. Urol. 7, 653–660 (2010).

3. Wagenlehner, F. et al. The Global Prevalence of Infections in Urology Study: A Long-Term, Worldwide Surveillance Study on Urological Infections. Pathogens 5, 10 (2016).

4. Geerlings, S. E. Clinical Presentations and Epidemiology of Urinary Tract Infections. Microbiol. Spectr. 4, 10.1128/microbiolspec.uti-0002–2012 (2016).

5. Hooton, T. M. Recurrent urinary tract infection in women. Int. J. Antimicrob. Agents 17, 259–268 (2001).

6. Saleem, Z. et al. Point prevalence surveys of health-care-associated infections: a systematic review. Pathog. Glob. Health 113, 191–205 (2019).

7. Timm, M. R., Russell, S. K. & Hultgren, S. J. Urinary tract infections: pathogenesis, host susceptibility and emerging therapeutics. Nat. Rev. Microbiol. 10.1038/s41579-024-01092-4 (2024) doi:10.1038/s41579-024-01092-4.

8. World Health Organization. Report on the Burden of Endemic Health Care-Associated Infection Worldwide. (World Health Organization, Geneva, 2011).

9. Flores-Mireles, A. L., Walker, J. N., Caparon, M. & Hultgren, S. J. Urinary tract infections: epidemiology, mechanisms of infection and treatment options. Nat. Rev. Microbiol. 13, 269–284 (2015).

10. Lacerda Mariano, L. & Ingersoll, M. A. The immune response to infection in the bladder. Nat. Rev. Urol. 17, 439–458 (2020).

11. Nielubowicz, G. R. & Mobley, H. L. T. Host–pathogen interactions in urinary tract infection. Nat. Rev. Urol. 7, 430–441 (2010).

12. Bryce, A. et al. Global prevalence of antibiotic resistance in paediatric urinary tract infections caused by *Escherichia coli* and association with routine use of antibiotics in primary care: systematic review and meta-analysis. BMJ i939 (2016) doi:10.1136/bmj.i939.

13. Simoni, A., Schwartz, L., Junquera, G. Y., Ching, C. B. & Spencer, J. D. Current and emerging strategies to curb antibiotic-resistant urinary tract infections. Nat. Rev. Urol. 10.1038/s41585-024-00877-9 (2024) doi:10.1038/s41585-024-00877-9.

14. Kao, J. S., Stucker, D. M., Warren, J. W. & Mobley, H. L. Pathogenicity island sequences of pyelonephritogenic Escherichia coli CFT073 are associated with virulent uropathogenic strains. Infect. Immun. 65, 2812–2820 (1997).

16. Rousset, F. et al. Genome-wide CRISPR-dCas9 screens in E. coli identify essential genes and phage host factors. PLOS Genet. 14, e1007749 (2018).

17. Wang, T. et al. Pooled CRISPR interference screening enables genome-scale functional genomics study in bacteria with superior performance. Nat. Commun. 9, 2475 (2018).

18. Rousset, F. & Bikard, D. CRISPR screens in the era of microbiomes. Curr. Opin. Microbiol. 57, 70– 77 (2020).

19. Barrangou, R. & Doudna, J. A. Applications of CRISPR technologies in research and beyond. Nat. Biotechnol. 34, 933–941 (2016).

20. Rousset, F. et al. The impact of genetic diversity on gene essentiality within the Escherichia coli species. Nat. Microbiol. 6, 301–312 (2021).

21. Baba, T. et al. Construction of *Escherichia coli* K-12 in-frame, single-gene knockout mutants: the Keio collection. Mol. Syst. Biol. 2, 2006.0008 (2006).

22. García, V. et al. Genome-wide analysis of fitness-factors in uropathogenic Escherichia coli during growth in laboratory media and during urinary tract infections. *Microb*. Genomics 7, (2021).

23. Shea, A. E. et al. Escherichia coli CFT073 Fitness Factors during Urinary Tract Infection: Identification Using an Ordered Transposon Library. Appl. Environ. Microbiol. 86, e00691–20 (2020).

24. García, V. et al. Genome-wide analysis of fitness factors in uropathogenic Escherichia coli in a pig urinary tract infection model. Microbiol. Res. 265, 127202 (2022).

25. Ma, J., Cai, X., Bao, Y., Yao, H. & Li, G. Uropathogenic Escherichia coli preferentially utilize metabolites in urine for nucleotide biosynthesis through salvage pathways. Int. J. Med. Microbiol. 308, 990–999 (2018).

26. Bouatra, S. et al. The Human Urine Metabolome. PLoS ONE 8, e73076 (2013).

27. Mann, R., Mediati, D. G., Duggin, I. G., Harry, E. J. & Bottomley, A. L. Metabolic Adaptations of Uropathogenic E. coli in the Urinary Tract. Front. Cell. Infect. Microbiol. 7, 241 (2017).

28. Bourne, C. R. Utility of the Biosynthetic Folate Pathway for Targets in Antimicrobial Discovery. Antibiotics 3, 1–28 (2014).

29. Yep, A., McQuade, T., Kirchhoff, P., Larsen, M. & Mobley, H. L. T. Inhibitors of TonB Function Identified by a High-Throughput Screen for Inhibitors of Iron Acquisition in Uropathogenic Escherichia coli CFT073. mBio 5, e01089–13 (2014).

30. Russo, T. A., Carlino, U. B. & Johnson, J. R. Identification of a New Iron-Regulated Virulence Gene, *ireA*, in an Extraintestinal Pathogenic Isolate of *Escherichia coli*. Infect. Immun. 69, 6209–6216 (2001).

31. King, J. E., Aal Owaif, H. A., Jia, J. & Roberts, I. S. Phenotypic Heterogeneity in Expression of the K1 Polysaccharide Capsule of Uropathogenic Escherichia coli and Downregulation of the Capsule Genes during Growth in Urine. Infect. Immun. 83, 2605–2613 (2015).

32. Umbarger, H. E. Biosynthesis of the Branched-Chain Amino Acids.

33. Kreisberg, J. F. et al. Growth Inhibition of Pathogenic Bacteria by Sulfonylurea Herbicides. Antimicrob. Agents Chemother. 57, 1513–1517 (2013).

34. Wu, J. & Gao, Y. Physiological conditions can be reflected in human urine proteome and metabolome. Expert Rev. Proteomics 12, 623–636 (2015).

35. Lawther, R. P. et al. Molecular basis of valine resistance in Escherichia coli K-12. Proc. Natl. Acad. Sci. 78, 922–925 (1981).

36. Jafari, N. V. & Rohn, J. L. An immunoresponsive three-dimensional urine-tolerant human urothelial model to study urinary tract infection. Front. Cell. Infect. Microbiol. 13, 1128132 (2023).

37. Flores, C. et al. A human urothelial microtissue model reveals shared colonization and survival strategies between uropathogens and commensals. Sci. Adv. 9, eadi9834 (2023).

38. LaRossa, R. A. & Schloss, J. V. The sulfonylurea herbicide sulfometuron methyl is an extremely potent and selective inhibitor of acetolactate synthase in Salmonella typhimurium. J. Biol. Chem. 259, 8753–8757 (1984).

39. Garcia, M. D. et al. Commercial AHAS-inhibiting herbicides are promising drug leads for the treatment of human fungal pathogenic infections. Proc. Natl. Acad. Sci. 115, (2018).

40. Hall, C. J., Mackie, E. R., Gendall, A. R., Perugini, M. A. & Soares da Costa, T. P. Review: amino acid biosynthesis as a target for herbicide development. Pest Manag. Sci. 76, 3896–3904 (2020).

41. Amorim Franco, T. M. & Blanchard, J. S. Bacterial Branched-Chain Amino Acid Biosynthesis: Structures, Mechanisms, and Drugability. Biochemistry 56, 5849–5865 (2017).

42. Brown, H. M. & Cotterman, J. C. Recent Advances in Sulfonylurea Herbicides. in Herbicides Inhibiting Branched-Chain Amino Acid Biosynthesis (ed. Stetter, J.) vol. 10 47–81 (Springer Berlin Heidelberg, Berlin, Heidelberg, 1994).

43. Zhou, Q., Liu, W., Zhang, Y. & Liu, K. K. Action mechanisms of acetolactate synthase-inhibiting herbicides. Pestic. Biochem. Physiol. 89, 89–96 (2007).

44. LaRossa, R. A. & Smulski, D. R. ilvB-encoded acetolactate synthase is resistant to the herbicide sulfometuron methyl. J. Bacteriol. 160, 391–394 (1984).

45. Duggleby, R. G., Pang, S. S., Yu, H. & Guddat, L. W. Systematic characterization of mutations in yeast acetohydroxyacid synthase: Interpretation of herbicide-resistance data. Eur. J. Biochem. 270, 2895–2904 (2003).

46. McCourt, J. A., Pang, S. S., Guddat, L. W. & Duggleby, R. G. Elucidating the Specificity of Binding of Sulfonylurea Herbicides to Acetohydroxyacid Synthase. Biochemistry 44, 2330–2338 (2005).

47. Varadi, M. et al. AlphaFold Protein Structure Database in 2024: providing structure coverage for over 214 million protein sequences. Nucleic Acids Res. 52, D368–D375 (2024).

48. Jumper, J. et al. Highly accurate protein structure prediction with AlphaFold. Nature 596, 583– 589 (2021).

49. Denamur, E., Clermont, O., Bonacorsi, S. & Gordon, D. The population genetics of pathogenic Escherichia coli. Nat. Rev. Microbiol. 19, 37–54 (2021).

50. De Bakker, V., Liu, X., Bravo, A. M. & Veening, J.-W. CRISPRi-seq for genome-wide fitness quantification in bacteria. Nat. Protoc. 17, 252–281 (2022).

51. van Opijnen, T. & Camilli, A. Transposon insertion sequencing: a new tool for systems-level analysis of microorganisms. Nat. Rev. Microbiol. 11, 435–442 (2013).

52. Liu, X. et al. Exploration of Bacterial Bottlenecks and *Streptococcus pneumoniae* Pathogenesis by CRISPRi-Seq. Cell Host Microbe 29, 107–120.e6 (2021).

53. Lamprecht, D. A. et al. Targeting de novo purine biosynthesis for tuberculosis treatment. Nature 644, 214–220 (2025).

54. Modi, G., Chacko, S. & Hedstrom, L. Targeting Purine Biosynthesis for Antibacterial Drug Design. 10.1039/9781782629870-00020(2017) doi:10.1039/9781782629870-00020.

55. Chua, S. M. & Fraser, J. A. Surveying purine biosynthesis across the domains of life unveils promising drug targets in pathogens. Immunol. Cell Biol. 98, 819–831 (2020).

56. Chan, C. C. Y. & Lewis, I. A. Role of metabolism in uropathogenic Escherichia coli. Trends Microbiol. 30, 1174–1204 (2022).

57. Hull, R. A. & Hull, S. I. Nutritional requirements for growth of uropathogenic Escherichia coli in human urine. Infect. Immun. 65, 1960–1961 (1997).

58. Vejborg, R. M. et al. Identification of Genes Important for Growth of Asymptomatic Bacteriuria Escherichia coli in Urine. Infect. Immun. 80, 3179–3188 (2012).

59. Mydock-McGrane, L. et al. Antivirulence C-Mannosides as Antibiotic-Sparing, Oral Therapeutics for Urinary Tract Infections. J. Med. Chem. 59, 9390–9408 (2016).

60. Flores, C. Bacterial adhesion strategies and countermeasures in urinary tract infection. Nat. Microbiol.

61. Sollier, J. et al. Revitalizing antibiotic discovery and development through in vitro modelling of in-patient conditions. Nat. Microbiol. 9, 1–3 (2024).

62. Calvo-Villamañán, A. et al. On-target activity predictions enable improved CRISPR–dCas9 screens in bacteria. Nucleic Acids Res. 48, e64–e64 (2020).

63. Rostain, W. et al. Cas9 off-target binding to the promoter of bacterial genes leads to silencing and toxicity. Nucleic Acids Res. 51, 3485–3496 (2023).

64. Mougiakos, I. et al. Efficient Genome Editing of a Facultative Thermophile Using Mesophilic spCas9. ACS Synth. Biol. 6, 849–861 (2017).

65. Wiktor, J., Lesterlin, C., Sherratt, D. J. & Dekker, C. CRISPR-mediated control of the bacterial initiation of replication. Nucleic Acids Res. 44, 3801–3810 (2016).

66. Love, M. I., Huber, W. & Anders, S. Moderated estimation of fold change and dispersion for RNA-seq data with DESeq2. Genome Biol. 15, 550 (2014).

67. Murphy, K. C. & Campellone, K. G. Lambda Red-mediated recombinogenic engineering of enterohemorrhagic and enteropathogenic E. coli. BMC Mol. Biol. 4, 11 (2003).

68. Bodenhofer, U., Bonatesta, E., Horejš-Kainrath, C. & Hochreiter, S. msa: an R package for multiple sequence alignment. Bioinformatics 31, 3997–3999 (2015).

69. Dereeper, A., et al. Phylogeny.fr: robust phylogenetic analysis for the non-specialist. Nucleic Acids Res. 36, W465–W469 (2008).

70. Letunic, I. & Bork, P. Interactive Tree of Life (iTOL) v6: recent updates to the phylogenetic tree display and annotation tool. Nucleic Acids Res. 52, W78–W82 (2024).

