## Extended Data Figures and supplementary results for "Genome-scale CRISPRi profiling reveals metabolic vulnerabilities of uropathogenic *Escherichia coli* in human urine"

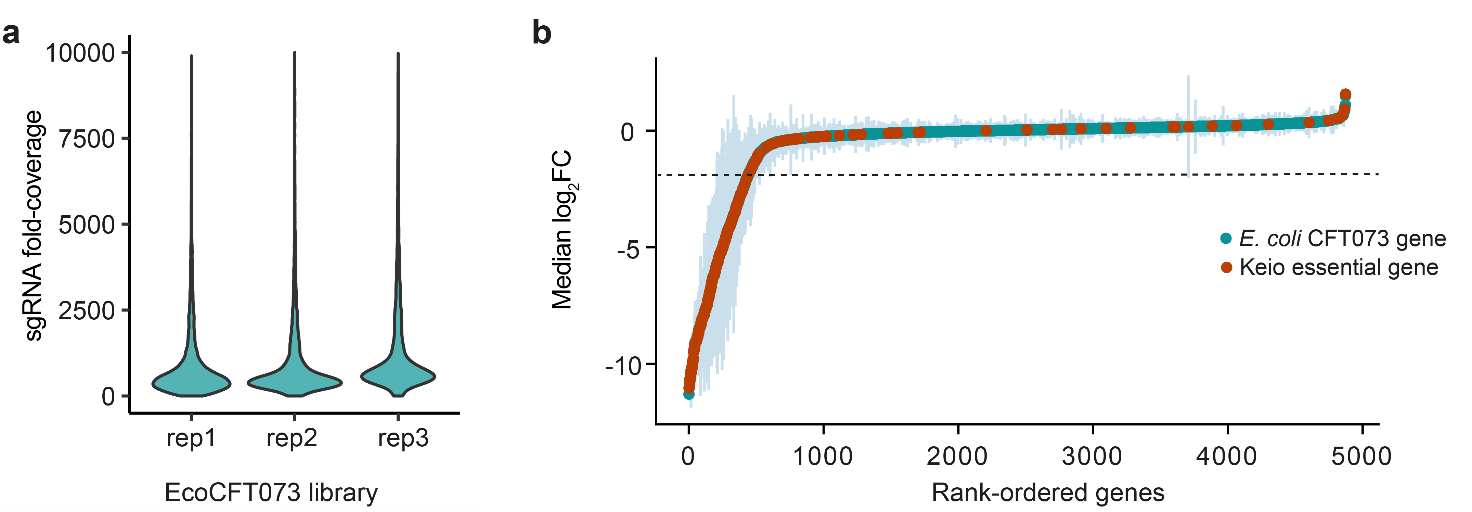

**Extended Data Figure 1: EcoCFT073 CRISPRi library generation and validation. a.** Distribution of sgRNA abundance across each EcoCFT073 CRISPRi library replicate. The average sgRNA fold coverage was 947, 937 and 1354 in library replicates 1, 2 and 3, respectively. **b.** Gene essentiality of *E. coli* CFT073 grown in LB medium. Genes were ranked by the median log_2_FC of all sgRNA targeting the same gene. Error bar represents the median absolute deviation (MAD). Keio essential genes are highlighted in red. The dashed line represents median log_2_FC of -2, the set threshold for essential genes.

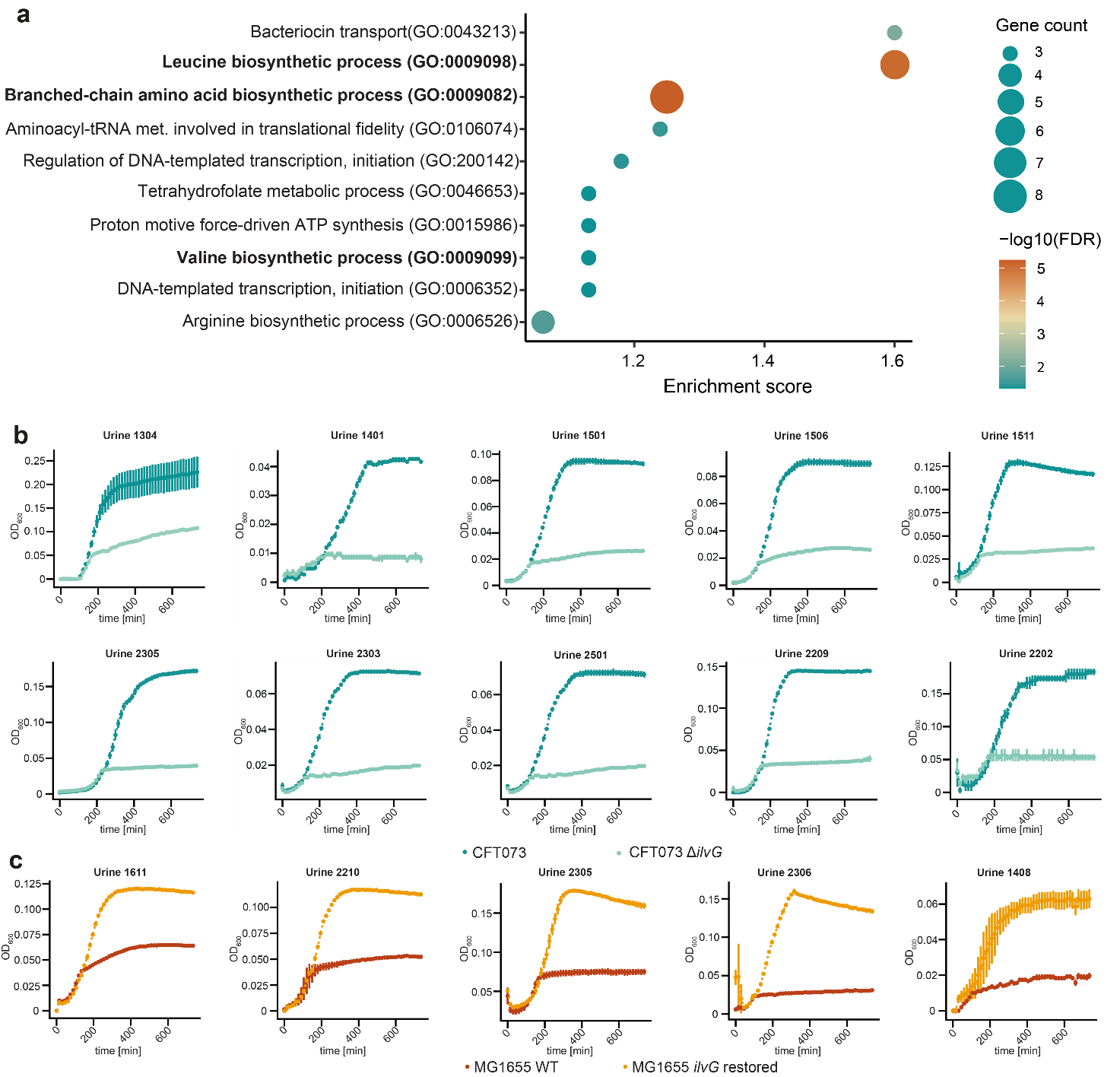

**Extended Data Figure 2: Validation of BCAA biosynthesis as conditionally essential in human urine. a.** Urine conditionally essential genes enrichment analysis using STRING. The dot plot shows the top 10 most significantly enriched Gene Ontology (GO) terms. Dot size corresponds to the number of genes involved in each pathway and dot color represents the -log10 false discovery rate (FDR). GO terms related to BCAA biosynthesis are highlighted in bold. **b.** Growth dynamics of *E. coli* CFT073 wild-type and Δ*ilvG* over 12 hours in urine samples donated by 10 healthy individuals**. c.** Growth dynamics of *E. coli* K12 MG1655 wild-type and *ilvG* restored over 12 hours in urine samples donated by 5 healthy people.

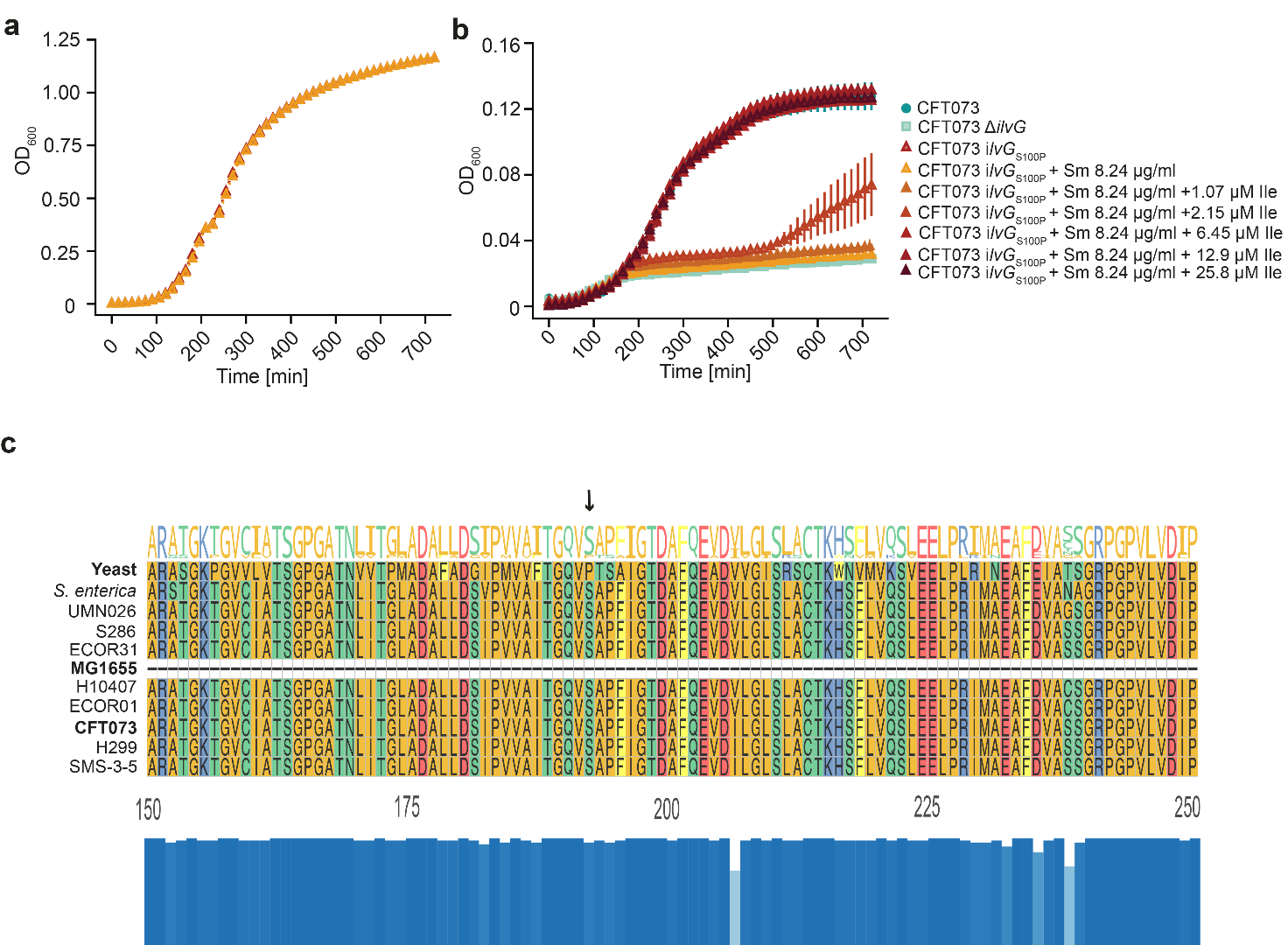

**Extended Data Figure 3: ALS II can be chemically targeted in UPEC. a.** Growth dynamics of *E. coli* CFT073 ilvG_S100P_ over 12 hours in MHB and in MHB supplemented with Sulfometuron-methyl (Sm) at 8.2 μg/ml (MIC measured in human urine). **b.** Growth dynamics of *E. coli* CFT073 *ilvG*_S100P_ over 12 hours in MHB supplemented with Sm (8.2 μg/ml) and increasing concentration of isoleucine (Ile). Grow dynamics of *E. coli* CFT073 wild-type and Δ*ilvG* are shown for comparison. Growth curves represent the mean ± standard deviation of three independent biological replicates. **c.** Multiple sequence alignment of ALS II proteins from different UPEC strains and *S. cerevisiae*. The figure shows amino acids 175-275 relative to the *S. cerevisiae* ALS II protein sequence. The bar char below the alignment indicates sequence conservation across 53 ALS II protein aligned. For clarity, the alignment figure displays only nine representative ALS II sequences from *E. coli* , while conservation is calculated from all. The black arrow mark residue 192 (relative to *S. cerevisiae* ALS II), which corresponds to proline in yeast and serine in all UPEC strain. This substitution (P192S) disrupts the interaction between ALS II and Sm.

**Supplementary results**

**Assessment of EcoCFT073 library reliability.** To evaluate EcoCFT073 library accuracy in predicting gene essentiality, we analyzed the fitness of Keio essential genes in LB. Out 300 Keio essential genes, 227 (75.6%) were also classified as essential in our screen, demonstrating the high reliability of EcoCFT073 library in predicting essential genes.

Among the remaining 73 Keio essential genes, 14 were either not annotated or absent in *E. coli* CFT073 genome (RefSeq entry: NZ_CP051263) and one is not targeted by any EcoCFT073 sgRNA (together representing ~5% of the Keio essential genes). The other 58 Keio essential genes (19.3%) showed no strong fitness defect in our LB screen, with median log2FC values ranging from 1.5 to -1.8. Different factors may explain these discrepancies. First, functional redundancy due to gene duplication or acquisition of genes with homologous function may compensate for the repression of the given target. For example, in the case of gene *grpE*, neither copy displayed a strong fitness defect, suggesting that the non-repressed copy compensated for the repressed one. In contrast, for genes such as *lptF*, *lptG* and *kdsB* one copy exhibited a strong fitness defect while the other one had little or no impact on *E. coli* CFT073 growth, implying that the second copy may be poorly expressed or inactive under the experimental condition tested. Finally, gene essentiality is also influenced by strain-genetic background. Notably, it was shown that *E. coli* K12, the strain underlying the Keio collection, exhibited the highest number of strain-specific essential genes, likely due to laboratory driven evolution.

**Comparison with previously performed genome-wide screens in human urine.** Our screen in pooled human urine showed overall good concordance with previously reported genome-wide studies at the functional level [ref], while also uncovering roughly two-thirds of genes not previously appreciated. Many of these novel candidates fall within pathways already implicated in fitness, whereas others broaden our understanding of additional processes required for growth in human urine (**Supp. Table 4 and 6**). At the same time, several genes previously classified as essential in other studies were not classified as such in our screen (**Supp. Table 4**).

These discrepancies can arise from several technical and analytical factors. First, differences in experimental design, including strain background, library saturation, screening conditions and urine source can substantially influence the fitness landscape. Second, distinct reference genomes were used to generate *E. coli CFT073* genome-wide libraries (RefeSeq entry CRISPRi library: NZ_CP051263, RefSeq entry transposons libraries: AE014075), which may have resulted in incomplete targeting or differences in gene annotation. Third, studies apply different criteria to define fitness or essential genes, which can lead to borderline cases being classified differently. For example, in two studies we used for comparison, *ilvB* was reported as fitness gene in human urine, in contrast with our findings. By deleting this gene, we confirmed that *ilvB* is not required for growth in human urine.
