## Supplementary material for "Genome-scale CRISPRi profiling reveals metabolic vulnerabilities of uropathogenic *Escherichia coli* in human urine": Python script used to generate EcoCFT073 CRISPRi library: EcoCFT073 add on library design.ipynb

```
# %% [code]
#Mapping EcoCG library on CFT073 complete genome
#Finding out the number of EcoCG sgRNAs that target a feature of CFCT073 genome

# %% [code]
#import ecoCG library
import pandas as pd

ecoCG=pd.read_csv("EcoCG library.txt", sep="\t", index_col=0)
ecoCG.head()


# %% [code]
#import the complete genome sequence of E.coli CFT073
from Bio import SeqIO

CFT=next(SeqIO.parse("CFT073.gb","genbank"))

# %% [code]
def revcomp(seq):
    trans=str.maketrans('ATGC','TACG')
    return seq.translate(trans)[::-1]

revcomp("ATGCGTCGA")

# %% [code]
#create a list of features at position (first create an empty list []). 
#Use the locus tag as unique feature qualifiers.

p=91282
def position_to_locustag(p):
    features_at_position=[]
    for feature in CFT.features[1:]:
        if p>=feature.location.start.position and p<=feature.location.end.position:
            locus_tag=feature.qualifiers["locus_tag"][0]
            if locus_tag not in features_at_position:
                features_at_position.append(locus_tag)
    
    return features_at_position

position_to_locustag(p)

# %% [code]
#re.finditer: it returns an object for every match it finds
#This dictionary gives the position of the guide from start to end and it excludes "edges guides" 
#(when PAM is locate inside the feature)

import re
from tqdm import tqdm

def get_guide_locus_tag(guide):
    tags=[]
    #forward orientation
    for match in re.finditer(guide+"[ATGC]GG", str(CFT.seq)):
        pam_pos=match.end()
        locus_tag=position_to_locustag(pam_pos)
        tags.append(locus_tag)

    
    #reverse orientation
    guide_revcomp=revcomp(guide)
    for match in re.finditer("CC[ATGC]"+guide_revcomp, str(CFT.seq)):
        pam_pos=match.start()
        locus_tag=position_to_locustag(pam_pos)
        tags.append(locus_tag)
        
    return tags

guide_tags_dict={}
for guide in tqdm(ecoCG.target.values):
    tags=get_guide_locus_tag(guide)
    guide_tags_dict[guide]=tags
        

# %% [code]
#to find out how many sgRNAs of EcoCG taraget a feature of CFT073 genome 
from collections import Counter
stats=Counter([len(tags) for tags in guide_tags_dict.values()])
for i in range(12):
    print("{} \tguides targeting {} features in CFT073".format(stats[i],i))

# %% [code]
#flatten the just generated list and find the loci target by the EcoCG library.
#So far we only know that 10906 ECoCG sgRNA targets one feature in CFT073 complete genome, 
# but we still don't know how many features are actually targeted by one guide 
#(EcoCG was designed to have 3 or 4 sgRNAs per gene)
def flatten(l):
    return [item for sublist in l for item in sublist]

loci_targeted_by_EcoCG=flatten(flatten(guide_tags_dict.values()))

# %% [code]
#To find out the number of CFT073 features targeted by EcoCG guides. EcoCG targets 3375 CFT073 features. 
# The remaining 1671 features has to be targeted by Add-on library sgRNAs. 
loci_targeted_by_EcoCG=set(loci_targeted_by_EcoCG)
len(loci_targeted_by_EcoCG)

# %% [code]
# Creation of add-on library 

# %% [code]
#Import CFT073 library

CFT_library=pd.read_csv("CFT073 library.csv",index_col=0)
CFT_library.head()

# %% [code]
#get information about the library: this library consists of 14942 sgRNAs
CFT_library.shape

# %% [code]
# The 14942 sgRNAs of the CFT073 library targets a total of 5022 features
len(CFT_library.locus_tag.unique())

# %% [code]
#Creation of the add-on library: 
#sgRNAs targeting features already targeted by EcoCG library are not required and therefore removed.
addon_library=CFT_library.loc[~CFT_library.locus_tag.isin(loci_targeted_by_EcoCG)]

# %% [code]
#Characteristics of add-on library: this novel library consists of 4877 guides. 
addon_library.shape

# %% [code]
# guides contained in the add-on library target a total 1649 features, while EcoCG sgRNAs targets 3375 CFT073 features
len(addon_library.locus_tag.unique()), len(loci_targeted_by_EcoCG)

# %% [code]
#Total number of features targeted by combining add-on library and EcoCG library: 5024. 
# CFT073 has a total of 5046 features. 22 features are not taergeted. 
#These leftover features are probably so small that don't caontain a PAM sequence and therefore a sgRNA cannot be created 
1649 + 3375

# %% [code]
#Add-on library inspection and export to file

# %% [code]
addon_library.head()

# %% [code]
#saving to file
addon_library.to_csv("addon_library_CFT073.csv")

# %% [code]
addon_library.guide

# %% [code]
#checking for the presence of control guides in EcoCG library
no_target_guides=[guide for guide in guide_tags_dict.keys() if len(guide_tags_dict[guide])==0]

# %% [code]
# 543 sgRNA does not target any CFT073 feature, meaning that they can be potentially used as control. 
# However non targeting guides that have a perfect identity in the seed region can cause off target effect
# therefore cannot be used as control 
len(no_target_guides)

# %% [code]
#excluding sgRNA with 9 bp perfect identity in the seed region: 
control_guides=[]
for guide in tqdm(no_target_guides):
    found=0
    #forward orientation
    for match in re.finditer(guide[-9:]+"[ATGC]GG", str(CFT.seq)):
        found=1
        
    #reverse orientation
    guide_revcomp=revcomp(guide)
    for match in re.finditer("CC[ATGC]"+guide_revcomp[:9], str(CFT.seq)):
        found=1
    
    if not found:
        control_guides.append(guide)

# %% [code]
# Out of 543 non targeting guides, only 64 can be used as control guide. 
# During data analysis, the Log2FC of the leftover 479 sgRNA has to be set to NaN, in order to exclude them from the analysis
len(control_guides)

# %% [code]
# to be even more precise, also sgRNA that match 8 bases, then mismatch and finally match again at least two bases in
# the seed region should not be used as control guide
#If alse these guides are excluded, a total of 52 sgRNA can be used as control guides: 
n=0
for guide in tqdm(control_guides):
    found=0
    #forward orientation
    for match in re.finditer(guide[-11:-9]+"[ATGC]"+guide[-8:]+"[ATGC]GG", str(CFT.seq)):
        found=1
        
    #reverse orientation
    guide_revcomp=revcomp(guide)
    for match in re.finditer("CC[ATGC]"+guide_revcomp[:8]+"[ATGC]"+guide_revcomp[9:11], str(CFT.seq)):
        found=1
        
    if not found:
        n+=1
        
print(n)

# %% [code]
#excluding sgRNA that match 8 bases, then mismatch and finally match again at least two bases in the seed region
#These are the control guides that are going to be used during data analysis
cont_guides = []
for guide in tqdm(control_guides):
    found=0
    #forward orientation
    for match in re.finditer(guide[-11:-9]+"[ATGC]"+guide[-8:]+"[ATGC]GG", str(CFT.seq)):
        found=1
        
    #reverse orientation
    guide_revcomp=revcomp(guide)
    for match in re.finditer("CC[ATGC]"+guide_revcomp[:8]+"[ATGC]"+guide_revcomp[9:11], str(CFT.seq)):
        found=1
        
    if not found: 
        cont_guides.append(guide)

# %% [code]
len(cont_guides)

# %% [code]
print(cont_guides)

# %% [code]
# export control guides (cont_guides) to an excel file
col1 = "sgRNA sequence"
data = pd.DataFrame({col1:cont_guides})
data.to_csv('control guides for CRISPRi screening.csv', index=False)

# %% [code]
# preparing a excel file containing sgRNAs that have to be excluded from data analysis 
# Those guides are: non targeting sgRNA except control guides and sgRNAs that target more than one CFT073 feature
# sgRNAs targeting 2 features
sgRNA_targeting_2_features =[guide for guide in guide_tags_dict.keys() if len(guide_tags_dict[guide])==2]
len(sgRNA_targeting_2_features)

# %% [code]
# sgRNA targeting 3 CFT073 features
sgRNA_targeting_3_features = [guide for guide in guide_tags_dict.keys() if len(guide_tags_dict[guide])==3]
len(sgRNA_targeting_3_features)

# %% [code]
# sgRNA targeting 4 CFT073 features
sgRNA_targeting_4_features = [guide for guide in guide_tags_dict.keys() if len(guide_tags_dict[guide])==4]
len(sgRNA_targeting_4_features)

# %% [code]
# sgRNA targeting 5 CFT073 features
sgRNA_targeting_5_features = [guide for guide in guide_tags_dict.keys() if len(guide_tags_dict[guide])==5]
len(sgRNA_targeting_5_features)

# %% [code]
# sgRNA targeting 6 CFT073 features
sgRNA_targeting_6_features = [guide for guide in guide_tags_dict.keys() if len(guide_tags_dict[guide])==6]
len(sgRNA_targeting_6_features)

# %% [code]
# sgRNA targeting 7 CFT073 features
sgRNA_targeting_7_features = [guide for guide in guide_tags_dict.keys() if len(guide_tags_dict[guide])==7]
len(sgRNA_targeting_7_features)

# %% [code]
# sgRNA targeting 9 CFT073 features
sgRNA_targeting_9_features = [guide for guide in guide_tags_dict.keys() if len(guide_tags_dict[guide])==9]
len(sgRNA_targeting_9_features)

# %% [code]
# sgRNA targeting 0 features
# cont_guides has to be excluded because they are actually needed for the analysis 
sgRNA_targeting_0_features = list(set(no_target_guides)-set(cont_guides))
len(sgRNA_targeting_0_features)

# %% [code]
# sgRNAs that have to be excluded from data analysis 
sgRNA_to_exclude = sgRNA_targeting_0_features +sgRNA_targeting_2_features + sgRNA_targeting_3_features + sgRNA_targeting_4_features + sgRNA_targeting_5_features + sgRNA_targeting_6_features + sgRNA_targeting_7_features + sgRNA_targeting_9_features  


len(sgRNA_to_exclude)

# %% [code]
#export list to an excel file
col1 = "sgRNA sequence"
data = pd.DataFrame({col1:sgRNA_to_exclude})
data.to_csv('sgRNAs that have to be excluded during data analysis.csv', index=False)

# %% [code]
# prepare an excel file containing only the guides that have to be considered from data analysis
ecoCG.to_csv("EcoCG library_MO.csv")
len(ecoCG)

# %% [markdown]
The just generated file contains all the EcoCG guides, including the one that have not to be considered during data analysis. Lateron, the Log2FC value of these not required guides it is going to be set to NaN.
During data analysis it is required an input file containing the sequence of all the guides. So I need to create a file containing both EcoCG guides as well as addo on guides. See below: 

# %% [code]
# Import EcoCG library and addon library 
import pandas as pd
addon = pd.read_csv("addon_library_CFT073.csv", index_col =0)
EcoCG = pd.read_csv("EcoCG library_MO.csv", index_col =0)
EcoCG

# %% [code]
addon.head()

# %% [code]
length_addon =len(addon)
print(length_addon)
length_EcoCG =len(EcoCG)
print(length_EcoCG)

# %% [code]
# Concatented addon sgRNAs with EcoCG library
addon_sgRNA= addon.guide
EcoCG_sgRNA = EcoCG.target
tot_sgRNA = [addon_sgRNA, EcoCG_sgRNA]
EcoCFT073 = pd.concat(tot_sgRNA)

# %% [code]
EcoCFT073

# %% [code]
#Convert EcoCFT073 (which contains all sgRNAs) into a pandas dataframe
col1 = "sgRNA"
EcoCFT073 = pd.DataFrame({col1:EcoCFT073})

# %% [code]
type(EcoCFT073)

# %% [code]
EcoCFT073

# %% [code]
# some sgRNA share the same sequence; those are removed from the list
# tot os sgRNA that were removed: 16508-16193 = 315
EcoCFT073 = EcoCFT073.drop_duplicates(keep = 'first')
EcoCFT073

# %% [code]
# Export the panda dataframe into a text file 
EcoCFT073.to_csv("EcoCFT073_no duplicates.txt", index= False, header = True)

#EcoCFT073.to_csv("EcoCFT073_v3.csv", index = True)

# %% [code]
EcoCFT073.to_excel("EcoCFT073_no duplicates.xlsx", index = False)

# %% [code]
prova = pd.read_excel("EcoCFT073_no duplicates.xlsx")
prova.head()

# %% [code]
# to run fast2q a csv file is necessary as imput, 
# therefore the created excel file containing sgRNA name and sequence is converted into csv
prova.to_csv("EcoCFT073_no duplicates_csv.csv", index = False)

# %% [markdown]
## Creation of EcoCFT073 library file with additional information

The library file created above contains only few information: the sgRNA sequence and an arbitrary sgRNA name. These characteristics are the only two required in order to run fast2q and therefore to generate a count table. However, later on, during data analysis, further information are required. It is for eaxample necessary to have know what is the gene targeted by the guide.

So, here below I will create a further EcoCF073 library file containing also info about the gene targeted by sgRNAs. 


# %% [code]
library = pd.concat(tot_sgRNA)
library

# %% [code]
#Convert library (which contains all sgRNAs) into a pandas dataframe
col1 = "target"
library = pd.DataFrame({col1:library})
library

# %% [code]
library.to_excel("EcoCFT073 with sgRNA sequences.xlsx", index = False)


# %% [code]
# in order to add a further colum  to the file, the two data frame have to share a column, 
# so using Excel I created a further coulmn containg sgRNA_name.
# the same column is going to be added to the dataframe containing info about gene (see below)
library =pd.read_excel("EcoCFT073 with sgRNA sequences.xlsx")
library

# %% [code]
# Extract genes info from both addon and EcCG library and concatenate then 
addon_gene = addon.gene
EcoCG_gene = EcoCG.gene_type
tot_gene = [addon_gene, EcoCG_gene]
gene = pd.concat(tot_gene)
gene

# %% [code]
# convert gene (which contains the target gene of each sgRNA) into a data frame
col1 = "gene"
gene_ID = pd.DataFrame({col1:gene})
gene_ID

# %% [code]
gene_ID.to_excel("EcoCFT073 with gene info.xlsx", index = True)

# %% [code]
gene_ID =pd.read_excel("EcoCFT073 with gene info.xlsx")
gene_ID

# %% [code]
# add a new column to library file, this column contains gene info
gene = gene_ID["gene"]

# %% [code]
library = library.join(gene)

# %% [code]
# remove sgRNA-name column from library dataframe 
library = library.drop(columns =['sgRNA_name'])
library

# %% [code]
# some sgRNA share the same sequence; those are removed from the list
# tot os sgRNA that were removed: 16508-16193 = 315
library = library.drop_duplicates(subset = ["target"],keep = 'first')
library

# %% [code]
# Export the created table into a text file 
numpy_array = library.to_numpy()
np.savetxt("EcoCFT073 library with sgRNA sequence and gene info.txt", numpy_array, fmt ="%s")

# %% [code]


# %% [code]


# %% [code]


# %% [code]


# %% [code]


# %% [code]


# %% [code]


# %% [code]
# to run fast2q a csv file is necessary as imput, 
# therefore the created excel file containing sgRNA name and sequence is converted into csv
prova.to_csv("EcoCFT073_no duplicates_csv.csv", index = False)

# %% [code]
# remove sgRNAs that share the same sequence from the list
 EcoCFT073_new = prova.drop_duplicates(subset =["ATAATGAGCCAGTATCTACT"], keep ="first")

# %% [code]
EcoCFT073=pd.read_csv("EcoCFT073_prova.txt", sep="\t", index_col=0)
EcoCFT073.head()

# %% [markdown]
## altro modo

# %% [code]
```
